# Regulation of Transcriptional Bursting and Spatial Patterning in Early *Drosophila* Embryo Development

**DOI:** 10.1101/2025.05.02.651973

**Authors:** César Nieto, Zahra Vahdat, Bomyi Lim, Abhyudai Singh

## Abstract

Nascent RNA synthesis often occurs in periods of high transcriptional activity, interspersed with basal or no activity periods. This phenomenon, known as transcriptional bursting, drives high intercellular variability in gene expression levels. How do key patterning genes in early *Drosophila melanogaster* embryos overcome this variability to establish precise spatial patterns for tissue development? To address this question, we study single cell transcriptional activity from MS2-based live imaging data of four transgenic constructs (*rho, Kr, sna* shadow, *sna* proximal enhancers) and the endogenous *eve* gene. We developed an algorithm to infer promoter states in hundreds of cells within the embryo using transcriptional activity data. Results show that while mean transcription levels exhibit spatial gradients, the burst duration and interburst timing remain surprisingly invariant across the embryo and different constructs. The time between consecutive bursts was consistent with a memoryless exponential distribution, whereas the burst duration exhibited tighter control with lesser stochastic variations. Our analysis identified two regulatory mechanisms for gene expression gradients: (1) similar burst-timing statistics across genes, with the activity time (the time from the first to the last burst) being modulated to regulate distinct expression levels; (2) for the same gene with different enhancers (*sna* shadow and *sna* proximal), we observed changes in the mean burst duration and variability of the inter-burst timing. This study provides a comprehensive approach to analyzing transcriptional bursting kinetics, revealing activity time as a major regulator of spatiotemporal expression patterning in early embryonic development.

## Introduction

Robust regulation of gene expression is essential for normal cell physiology and embryo development. Variations in expression levels or ectopic activation at inappropriate times or locations can significantly affect cell phenotypes, fitness, survival, and differentiation [1–3]. Gene misregulation can be related to different developmental defects and disease phenotypes [4, 5]. Despite the need for strict control, single cell imaging has revealed random fluctuations (noise) in promoter transcriptional activity over time [6, 7]. Several studies have characterized transcriptional noise in the expression of developmental genes across organisms [8–15], and it is important to understand how this noise in single cells affects overall spatial expression patterning and subsequent development.

Gene expression is characterized by transcriptional bursts: intermittent periods of promoter activity followed by periods of inactivity in which transcripts are not produced [16–26]. Randomness in switching between active and inactive promoter states has been shown to be a key driver of intercellular variability in gene product levels across isogenic cell populations [27–37]. By analyzing the properties of transcriptional bursting in various genetic backgrounds, we can better understand the mechanisms for regulating the levels of gene products [16, 38, 39].

We present a comprehensive analysis of the transcriptional bursting properties of reporter genes driven by various regulatory enhancers and endogenous genes in early *Drosophila* embryos. We analyze publicly available live imaging data where single-cell transcriptional activity is tracked as a function of time and position in the embryo [40,41]. Each gene is labeled with the 24x MS2 sequence, which allows visualization of nascent transcripts in each cell with a fluorescent marker (MCP-GFP), allowing high temporal resolution to investigate the dynamics of bursting [42, 43]. We develop a simple yet robust algorithm to estimate promoter transcriptional states based on the accumulation and decay of individual fluorescence trajectories. These inferred promoter states are then used to quantify the spatiotemporal statistics of transcriptional bursting across the gene expression domain.

Among the properties of transcriptional bursting studied, we consider the burst duration (ON time), the time between successive bursts (OFF time), the rate of nascent mRNA transcription (loading rate), and the span from the first to the last burst (activity time). To examine how parameters such as spatial heterogeneity, different genes, or different enhancers affect burst dynamics, we selected enhancers of *Krüppel (Kr)* and *rhomboid (rho)* genes. The mean transcriptional activity of these genes drives expression that follows a graded pattern along the anterior-posterior and dorsoventral axes of an embryo, respectively [44, 45]. Interestingly, we find that the mean ON and OFF times are homogeneous across the expression domain, not showing the expression gradient observed in the average transcription activity. This heterogeneity can be mainly explained by differences in activity time, suggesting that different cells have different durations of active transcription but similar bursting properties once activated.

We compare two redundant *sna* enhancers (*sna* shadow and *sna* proximal), which drive spatially homogeneous ventral expression, revealing distinct bursting characteristics and indicating that enhancers can modulate burst timing statistics [46]. Furthermore, we study the endogenous *even-skipped* (*eve*) gene, with its seven-strip spatial pattern [47, 48]. Although *eve* shows homogeneous burst duration, we also found spatially varied activity time and inter-burst duration, aligning with observations from the *Kr* and *rho* constructs.

## Results

This study characterizes the spatiotemporal dynamics of key genes involved in *Drosophila* embryonic development. We analyze gene expression dynamics during the 14th nuclear cycle (NC14), which corresponds approximately to a time window of 2-3 hours after fertilization. We use the enhancers of *rho, Kr*, and *sna*, which drive transcription of the MS2 reporter genes that recapitulate endogenous gene expression [21]. We also analyze the transcription of the endogenous *eve* gene (see Methods). All of these genes play an important role in the formation of body patterns in *Drosophila* [49].

### Spatio-temporal properties of transcription bursts

The experiments analyzed (see Methods) consist of confocal microscope images of living *Drosophila* embryos (Figure 1A). These images discern fluorescence dynamics from individual cells, allowing single-cell resolution analysis along the anteriorposterior and dorsoventral axes of the embryo (Figure 1B). This fluorescence trajectory corresponds to transcription dynamics captured using the MS2/MCP system (Figure 1C). In this system, the number of nascent mRNA molecules is proportional to the fluorescence signal (Figure 1D, top panel). This trajectory reveals different periods of intense activity (referred to as bursts) with stochastic duration indicated by *τ*_*ON*_ and intervals between successive bursts denoted as *τ*_*OFF*_ . To estimate the transcriptional state, we developed an algorithm (see Methods) based on the finding that the fluorescence signal increase (during ON state) and decrease (during OFF state) is approximately linear over time (Figure S1), with relatively similar slopes (Figure S2). In Figure 1D bottom, we show an example of an inferred transcription state trajectory from the fluorescence signal (see Supplementary Information Section S3 and Figure S3 for technical details of the inference algorithm).

**Figure 1:**
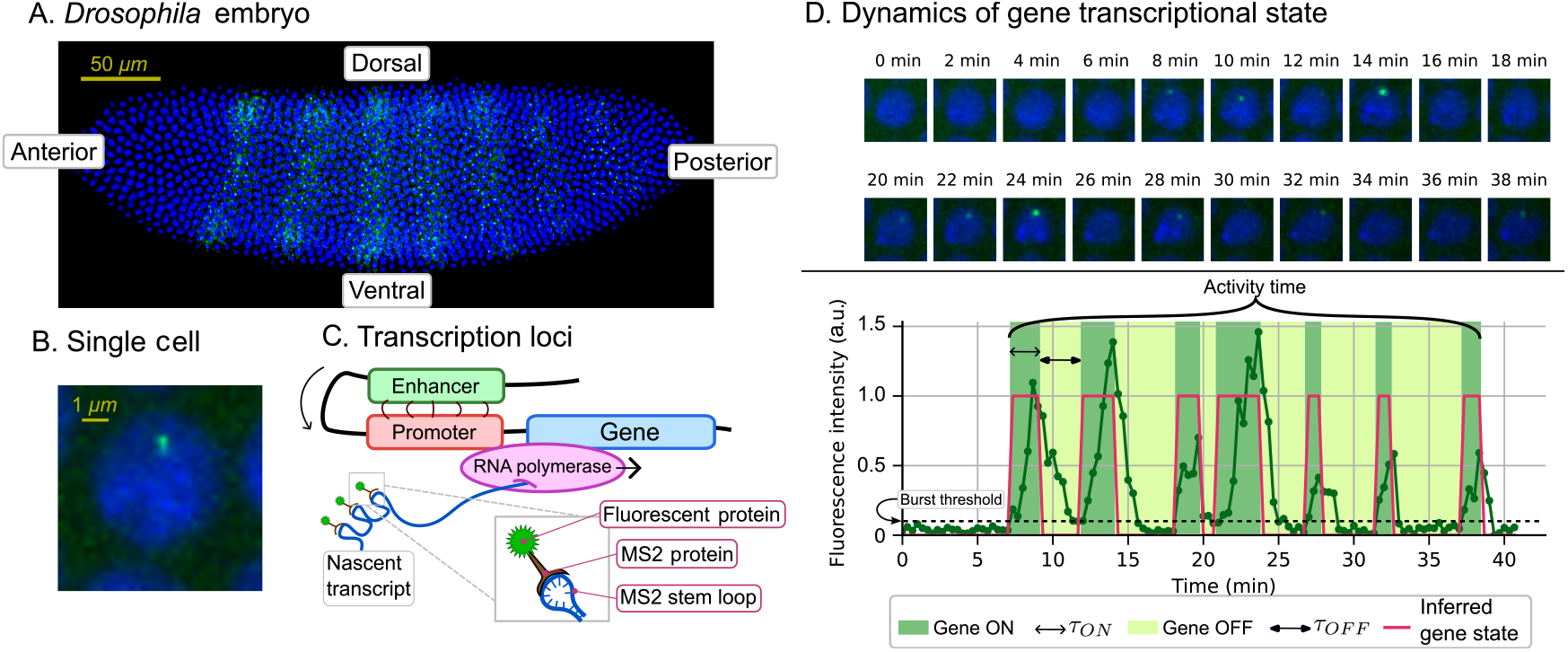
The MCP/MS2-mediated live imaging system enables the visualization of transcriptional activity at the single-cell level in living *Drosophila* embryos. **(A)** Representative snapshot of MS2-based imaging, where endogenous *eve* is tagged with MS2 to visualize the spatial expression pattern in embryos. Regions of the embryo (ventral, dorsal, anterior and posterior) are provided. **(B)** A single cell in the embryo showing GFP fluorescence with intensity assumed to be proportional to the number of nascent mRNAs. **(C)** Schematic showing how the nascent transcripts are tagged with fluorescent proteins using the MS2/MCP system. **(D)** (Top:) Representative snapshots of a single nucleus producing transcripts (green puncta) over time. (Bottom:) Fluorescence dynamics over time (green line) and the inferred transcriptional state (red line). From the inferred state, we estimate the duration of active (ON) states (*τ*_*ON*_ ) and the duration of inactive (OFF) states (*τ*_*OFF*_ ). The black dashed line shows the burst threshold: the level where any sudden increase from the base level is considered as a burst (see Methods). Bracket in the top of the trajectory represents the activity time: the time between the beginning of the first burst and the end of the last one.

The fluorescence intensity averaged throughout the nuclear cycle reveals a distinct spatial pattern in the embryo (Figure 1A). Here, we relate this mean fluorescence to the statistics of *τ*_*ON*_ and *τ*_*OFF*_ . In addition to *τ*_*ON*_ and *τ*_*OFF*_ , our algorithm also estimates the *loading rate λ*^*^, which is defined as the signal increase rate that best predicts the signal dynamics (see Methods) and is specific for each cell. Finally, we also estimate the time between the beginning of the first burst and the end of the last burst, which is called the *activity time*. We will study how these different burst properties vary spatially across the embryo, and which properties have the best concordance with the single-cell averaged mean fluorescence level.

### The activity time as an important contributor to the control of gene expression

We first analyze the burst statistics of the embryos expressing *rho* and *Kr* genes. *rho* shows a graded expression along the dorsoventral (vertical) axis and *Kr* shows a gradient along the anterior-posterior (horizontal) axis (see Figure 2A and 2G) [44, 45]. *rhoNEE* enhancer that drives the neuroectodermal expression of *rho* is used to drive the reporter gene *MS2-yellow* expression [45]. On the other hand, the expression pattern of *Kr* is driven by the enhancer *Kr CD2* [50].

**Figure 2:**
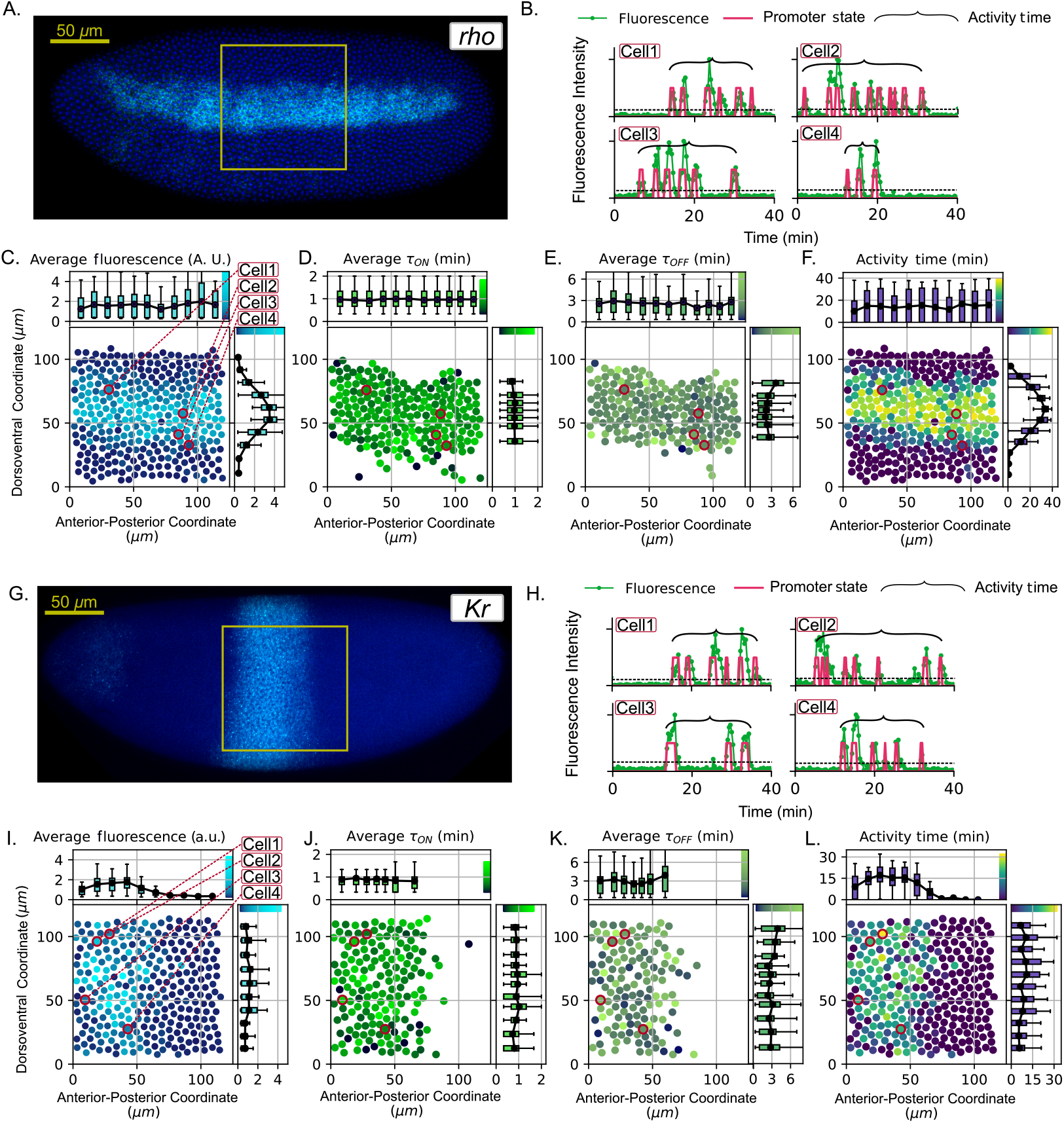
Heterogeneous spatial expression patterns for enhancer *rho* and *Kr*. (**A**,**G**) Gene expression driven by the *rho* (A) and *Kr* (G) enhancers which exhibit a dorsoventral and anterior-posterior gradient, respectively. The yellow box indicates, approximately, the region imaged by live imaging. (**B**,**H**) Representative fluorescence trajectories (green) and the inferred transcriptional state (red) for *rho* (B) and *Kr* (G). Fluorescence signals are normalized to their maximum. Activity time is represented by black brackets. Dashed horizontal line represents the burst threshold. (**C**,**I**) Average fluorescence intensity, (**D**,**J**) cell averaged *τ*_*ON*_ , (**E**,**K**) average *τ*_*OFF*_ , and (**F**,**L**) the activity time. The position of each point represents one cell in the beginning of NC14 and its color represents the mentioned average burst property. Cells presented in (B,H) are highlighted in red in (C-F) and (I-L) respectively. Marginal plots show the mean variables calculated with a spatial binning. The error bars show the 5% and 95% quantiles, while the length of the boxes show the 25% and 75% quantiles for each bin. Black dots represent the mean value. The means are connected by lines to show the spatial trends. Marginal plots also present the color scale at their edges used in the central scatter plot.

Our initial hypothesis was that gene expression can be directly associated with the frequency and duration of transcriptional bursts. This means that in regions with low or high mean gene expression, we expect less or more frequent bursts, respectively. To verify the validity of this hypothesis, we plot the spatial patterns of the average fluorescence intensity of *rho* (Figure 2C-F). As examples of observed dynamics, four representative nuclei from the gene expression domain were selected and their transcriptional trajectories were plotted (Figure 2B). In Figure 2C, we show a scatter plot with each dot representing one cell. The position of each dot represents the position of the respective tracked cell at the beginning of NC14. The color of the dots represents the fluorescence level averaged throughout NC14. To better visualize the spatial gradient of the burst properties, we also made marginal plots by grouping the cells into bins along the dorsoventral and anterior-posterior axis and calculating the statistics for each bin.

Figure 2D shows spatial patterns of the cell-averaged *τ*_*ON*_ . We excluded cells that did not show fluorescence levels above the burst threshold, as it is not possible to estimate the average *τ*_*ON*_ . The marginal plots in Figure 2D show the variation of the cellaveraged *τ*_*ON*_ for the selected bins. These bins reveal that the mean *τ*_*ON*_ for multiple regions of the embryo is constant, with a value of about 1 min. Similar spatial trends for *τ*_*OFF*_ are presented in Figure 2E. Marginal plots show that the mean *τ*_*OFF*_ along the axes also shows a low variation throughout the embryo. In this case, the average *τ*_*OFF*_ is around 3 min.

The finding that the cell averaged *τ*_*ON*_ and *τ*_*OFF*_ is spatially uniform contradicts the hypothesis of a non-homogeneous gene switching rate. To explain pattern formation in the selected genes, we noticed, as presented in Figure 2B, that the dynamics of gene expression can be simplified as periods of activity. During these long periods, multiple bursts of similar duration occur at an approximately constant rate. This motivated us to define the activity time as the span from the first burst to the final one. The duration of this activity time is marked with a bracket on the signal trajectory in Figure 2B.

We notice that spatial gene expression corresponds better to the patterns of activity time (Figure 2F) than to the cell-averaged *τ*_*ON*_ and *τ*_*OFF*_ . This suggests that the activity time better reflects the spatial heterogeneity of gene expression. This is confirmed by the *Kr* construct, where mean fluorescence shows a similar spatial pattern to activity time (Figures 2G-2L). Although *Kr* has different mean fluorescence levels along the embryo axis compared to *rho*, the average *τ*_*ON*_ and *τ*_*OFF*_ in *Kr* are relatively homogeneous with mean durations similar to those in *rho*. Our analysis indicates that the graded expression pattern results from the modulation of activity time rather than bursting properties.

### Different enhancers can modify the time between successive bursts

Enhancers are short non-coding DNA sequences that interact with the target promoter to drive transcription (Figure 1C). Particular enhancers can increase the likelihood of transcriptional initiation by containing binding sites for transcription factors and other proteins that promote transcription [51]. We now ask whether different enhancers of the same gene can affect the burst-timing statistics. We studied the *sna* gene, which has a broad expression domain on the ventral side of an embryo, as shown in Figure 3A. *sna* is regulated by two enhancers, the proximal enhancer (*snaPE* ) that is located right upstream of the transcription start site (TSS) and the distal or shadow enhancer (*snaSE* ) located 8kb upstream of the TSS [46]. Both enhancers are known to drive *sna* expression in the same domain at the same time, but *snaPE* drives weaker expression than *snaSE* (Figures 3B and 3G) [52].

**Figure 3:**
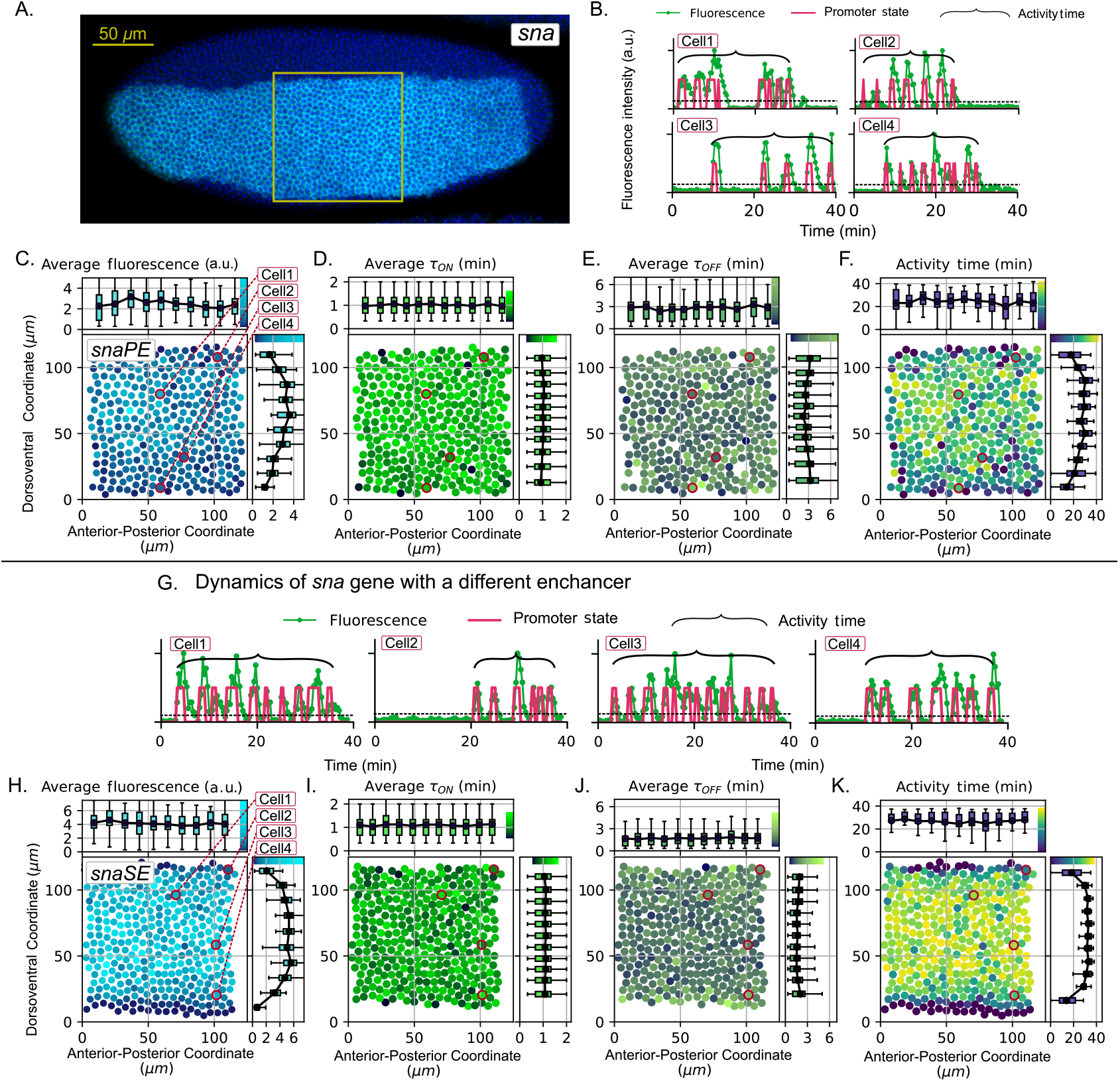
Spatial patterns of *sna* gene expression for different enhancers: shadow (SE) and proximal (PE) (**A**) Image of the *sna* expression across the embryo. The yellow box indicates, approximately, the region imaged. (**B, G**) Representative gene expression trajectories normalized to the maximum (green) and the inferred transcriptional state (red) for *sna*PE (B) and *sna*SE (G). Activity time is represented by black brackets. Dashed horizontal line represents the burst threshold. These cells are highlighted in red in (C-F) and (H-K). (**C, H**) Spatial trends of average fluorescence, (**D, I**) average *τ*_*ON*_ , (**E, J**) average *τ*_*OFF*_ , and (**F, L**) activity time. Marginal plots show how the mean changes in different spatial bins of the expression domain. Bar length represents the 25-75% interquantile distance and error bars, the 5-95% interquantile distance.

Comparing the bursting properties of *snaPE* and *snaSE*, we find some similarities with the *rho* and *Kr* genes. For example, *τ*_*ON*_ and *τ*_*OFF*_ have low spatial heterogeneity and the activity time is highly correlated with the average fluorescence (Figures 3C-F and 3H-K). However, some differences are also observed. For example, the mean fluorescence intensity is higher for the SE enhancer than for the PE enhancer, although both regulate the same gene (Figures 3C and 3H). This difference can be explained by a shorter average *τ*_*OFF*_ for the SE enhancer. Since *snaSE* is considered a stronger enhancer than *snaPE*, producing more mRNA at a given duration, our results suggest that *snaSE* can drive stronger expression by spending less time in the OFF state, which can also explain similar results found previously [40]. Our analysis indicates that different enhancers of the same gene can induce different bursting behaviors and therefore regulate the level of mRNA production differently.

### Statistical analysis of transcriptional bursting events

To better understand the statistical properties of the burst timing variables, we collect all the values of *τ*_*ON*_ and *τ*_*OFF*_ for different replicate embryos and study their statistical distribution without discrimination by cell or position. More specifically, we quantify the mean *τ*_*ON*_ and *τ*_*OFF*_ (denoted by ⟨*τ*_*ON*_⟩ and ⟨*τ*_*OFF*_⟩, respectively). The random variability in *τ*_*ON*_ and *τ*_*OFF*_ is measured by the squared coefficient of variation (the variance over the mean squared) and represented by 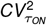 and 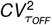, respectively. These statistics are estimated over all cells in the embryo during NC14 for 3 different embryos (biological replicas) per construct.

### While *τ*_*OFF*_ follows an exponential distribution, *τ*_*ON*_ has a hypo-exponential distribution

The analysis of ⟨*τ*_*ON*_ ⟩ and ⟨*τ*_*OFF*_ ⟩ is presented in Figure 4A. There, we show that ⟨*τ*_*ON*_ ⟩ ≈ 1 min while ⟨*τ*_*OFF*_ ⟩ ≈ 3 min for *Kr, rho*, and *snaPE*. The main outlier corresponds to *snaSE* ⟨*τ*_*OFF*_⟩ ≈ 2 min. These results show that the properties of the burst dynamics are consistent not only for different replicas, but also among different genes. Regarding the random variability of the time intervals, Figure 4B shows how while 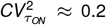 in all replicas and genes, 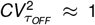 for *Kr, rho*, and *snaPE*. The main outlier, *snaSE*, shows a variability of 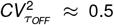. As a reference for the interpretation of this metric, we recall that the coefficient of variation for an exponentially-distributed random variation is 1 (dashed line in Figure 4B).

**Figure 4:**
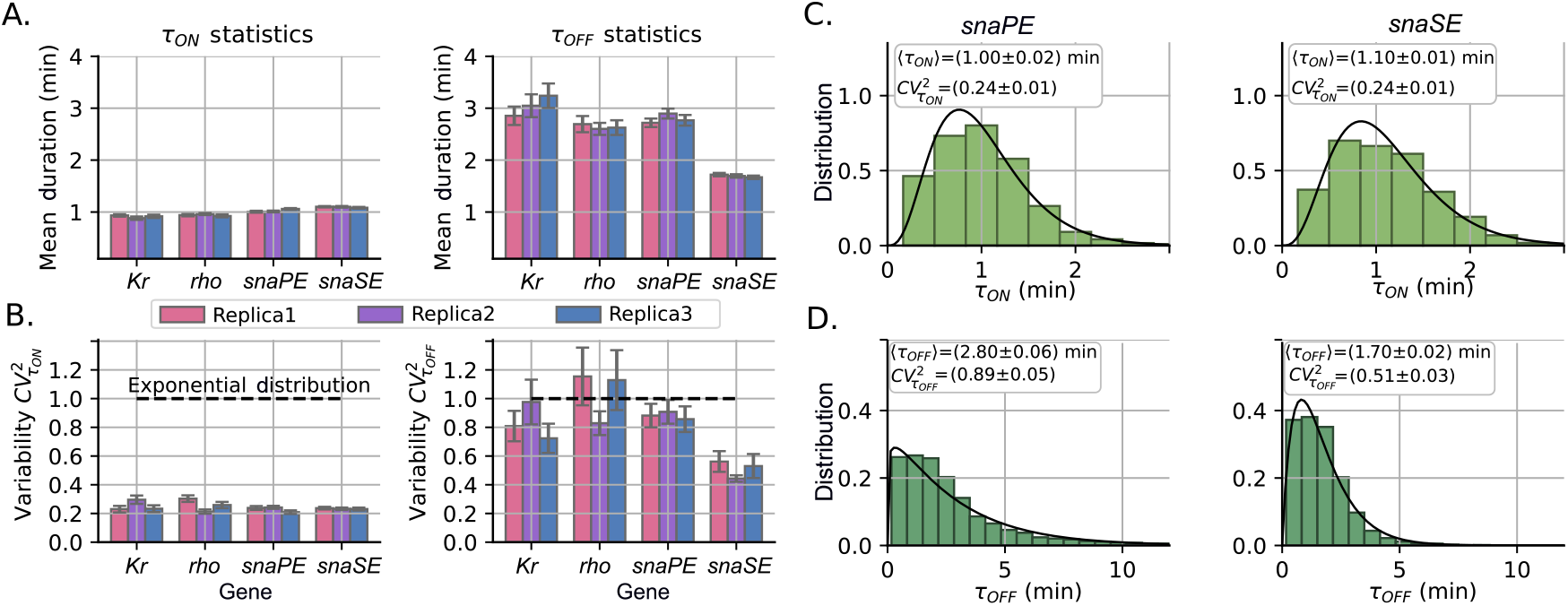
Statistical properties of transcriptional bursting timing across different constructs. (**A**) Mean duration of *τ*_*ON*_ and *τ*_*OFF*_ for the four studied enhancers, each with three replicas. (**B**) Random variability of *τ*_*ON*_ and *τ*_*OFF*_ measured by the squared coefficient of variation 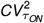 and 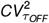 respectively. The dashed line represents the coefficient of variation of one for an exponentially-distributed random variable. (**C**) Probability density for *τ*_*ON*_ for the *sna* enhancers. The PE enhancer (left) is compared to the SE enhancer (right). The mean and squared coefficients of variation are presented with the 95% confidence interval. The solid line represents the best fit to a gamma distribution. (**D**) Analysis similar to (C) but for *τ*_*OFF*_ . The resolution of *τ*_*ON*_ and *τ*_*OFF*_ is 20 s.

To better visualize the interpretation of the metrics of 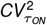 and 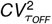, we plot the histograms of *τ*_*ON*_ for *snaPE* and *snaSE* (Figure 4C). For these enhancers 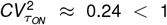. This means that the distribution of *τ*_*ON*_ is hypoexponential, i.e., with less random variation than an exponential distribution (which has a coefficient of variation of 1). We also plot the histogram of *τ*_*OFF*_ in Figure 4D for the same enhancers. *τ*_*OFF*_ for *snaPE* has a mean of approximately 2.8 min and a noise of 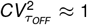 similar to the exponential distribution. *snaSE*, on the contrary, shows a shorter ⟨*τ*_*OFF*_ ⟩ ≈ 1.7 min and a lower variability 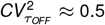.

### Correlation between burst variables an mean fluorescence

In this section, we explicitly examine how the cell-averaged fluorescence relates to four other variables: cell-averaged *τ*_*ON*_ , cellaveraged *τ*_*OFF*_ , activity time and optimal loading rate *λ*^*^ (equation (3) in Methods). We gather data from three biological replicate embryos for each construct and observe the trends between these variables.

We observe that activity time increases monotonically with increasing mean fluorescence intensity, reaching saturation at high levels of mean fluorescence (Figure 5A). On the other hand, cellaveraged *τ*_*ON*_ , cell-averaged *τ*_*OFF*_ , and *λ*^*^ show weaker correlations with mean fluorescence. Cell-averaged *τ*_*ON*_ increases slightly with increasing mean fluorescence, validating our simplification that *τ*_*ON*_ is essentially constant across transgenic constructs (Figure 5B). The cell-averaged *τ*_*OFF*_ shows a slightly decreasing trend with mean fluorescence, although the trend is weaker compared to the relationship between activity time and mean fluorescence (Figure 5C). Relatively longer periods of *τ*_*OFF*_ are observed for nuclei with lower gene expression. Finally, there is almost no correlation between the loading rate *λ*^*^ and the mean fluorescence (Figure 5D).

**Figure 5:**
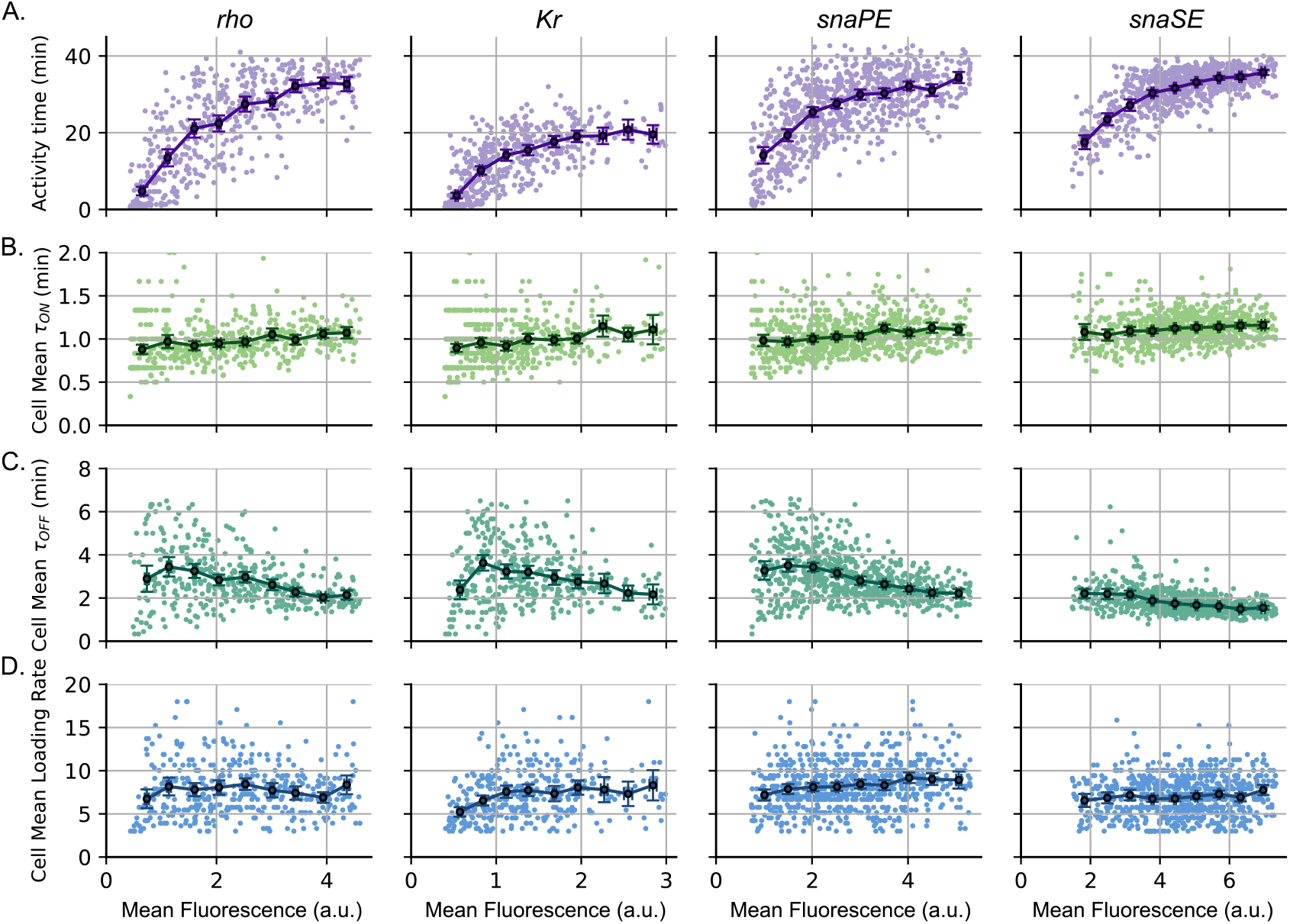
Statistical properties of the transcriptional bursts with respect to the mean fluorescence for different enhancers. We plot the trend of different signal statistics such as (**A**) activity time, (**B**) mean ON state duration, (**C**) mean OFF state duration, and (**D**) mean loading rate (the rate of signal increase) relative to the mean fluorescence. Each dot in the plots represents the mean value of a single cell obtained from its fluorescence trajectory during NC14. These trends suggest that the activity time aligns better with mean fluorescence.

### The statistics of the *eve* gene expression

Until now, the data analyzed to estimate the burst statistics correspond to transgenic constructs. Next, we performed a similar analysis of the transcriptional burst statistics of *eve*, an endogenous gene with a more complex spatial pattern of expression. The average fluorescence level of *eve* has a pattern of seven stripes during early embryonic development (Figure 6A) [47]. These stripes are the result of interactions between *eve* and gap genes defining the position of specific para-segments that set the stages for the final segmentation of the embryo body [53]. As observed in Figure 6A, we analyze transcription trajectories of the nuclei corresponding to stripes 2-5. Examples of typical fluorescence trajectories of *eve* expression can be seen in Figure 6B showing, in general, a lower average intensity than the synthetic constructs (*Kr, rho* and *sna*) but a similar basal intensity. Regarding the timing statistics, we observe low spatial variability on the cell-averaged *τ*_*ON*_ and *τ*_*OFF*_ on the spatial domain of *eve* (Figure 6D and 6E). Furthermore, these mean times also show values similar to those of the synthetic constructs: ⟨ *τ*_*ON*_ ⟩ ≈ 1 min and ⟨*τ*_*OFF*_⟩ ≈ 3 min. Again, the activity time appears to align better with the average fluorescence pattern (Figure 6F).

**Figure 6:**
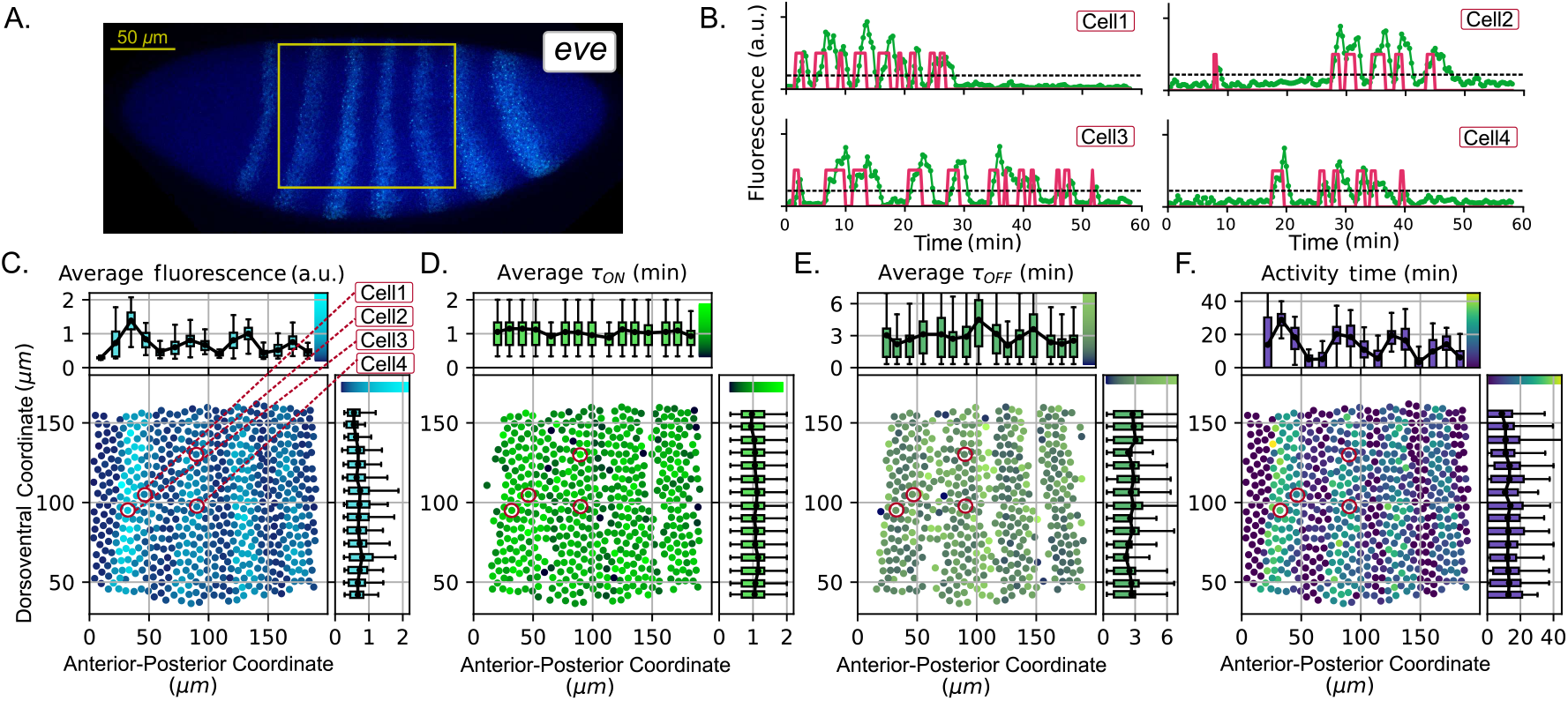
Spatial patterns of endogenous *eve* gene expression. (**A**) Expression pattern of the *eve* gene. The yellow box indicates, approximately, the region imaged. (**B**) Expression trajectories normalized to the maximum (green) and the state of the inferred gene (red). Activity time is represented by black brackets. The horizontal dashed line represents the burst threshold which is the same as used in the constructs. These cells are highlighted in red in (C-F). (**C**) Spatial trends of average fluorescence, (**D**) average *τ*_*ON*_ , (**E**) average *τ*_*OFF*_ , and (**F**) the activity time in the studied embryo domain. Bar length represents the 25-75% interquantile distance and error bars, the 5-95% interquantile distance.

Figure 7 presents a more detailed analysis of the relationship between mean fluorescence and burst properties for the *eve* gene, based on data from six biological replicates. As with synthetic constructs, cell-averaged fluorescence shows a strong alignment with activity time (Figure 7A), but only a weak correlation with cell-averaged *τ*_*ON*_ (Figure 7B) and loading rate (Figure 7D). Cell-averaged *τ*_*OFF*_ shows a stronger decrease with mean fluorescence compared to the one observed in synthetic constructs (Figure 7C). The mean burst duration for *eve* shows relative consistency over replicas with similar mean (Figure 7E) and 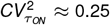 (Figure 7F). However, *τ*_*OFF*_ has slightly less consistency on the mean ⟨*τ*_*OFF*_ ⟩ while the random variability 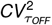 was hyperexponential 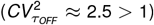.

**Figure 7:**
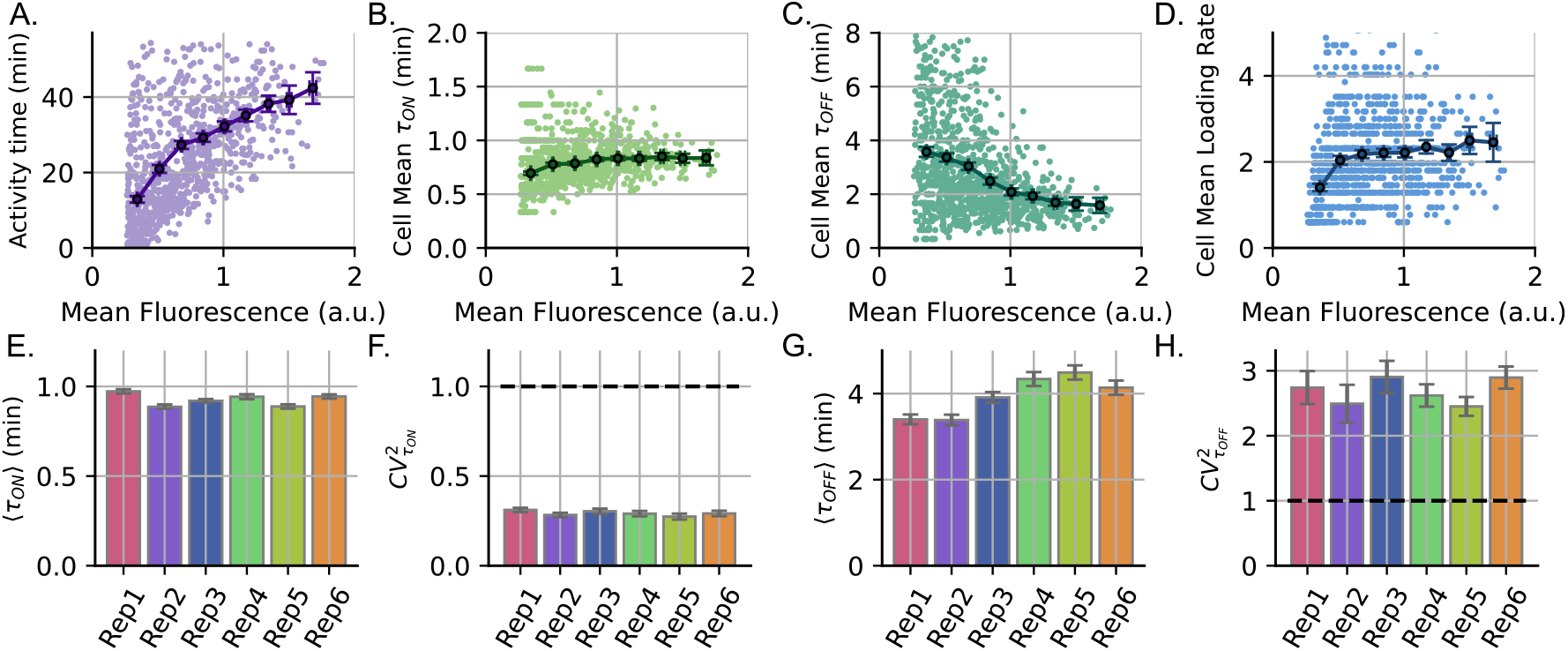
Properties of transcriptional burst timing for the expression of endogenous *eve* gene. Plots of average burst properties with respect to single-cell averaged fluorescence level (**A**) activity time. (**B**) Mean *τ*_*ON*_ , (**C**) Mean *τ*_*OFF*_ and (**D**) Mean loading rate. Each dot in these plots corresponds to a cell and the statistics are calculated over the NC14 time-span. Error bars represent the binned mean trend with 95% confidence. We also present the statistics of the burst timing estimated using all bursts and all cells for 6 different biological replicas: (**E**) mean burst duration ⟨*τ*_*ON*_ ⟩, (**F**) variability on burst duration 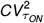, (**G**) mean time between successive bursts ⟨*τ*_*OFF*_ ⟩ and (**H**) variability in the time between bursts 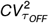. Horizontal dashed lines in (F) and (H) correspond to a coefficient of variation of 1 for the exponential distribution. Error bars represent the 95% confidence interval.

The relatively large 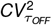 observed in *eve* is mainly due to the presence of abnormally long inter-burst times, often exceeding 12 min (Figure S4A). To validate this concept, we reanalyzed the burst statistics, focusing exclusively on cells with inter-burst times of less than 12 min. After excluding these cells, all burst properties align with those observed in synthetic constructs: ⟨*τ*_*ON*_ ⟩ ≈ 1 min and ⟨*τ*_*OFF*_ ⟩ ≈ 3 min, while 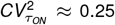 and 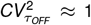 (Figure S4C-D). The comparison of the burst properties versus the mean fluorescence after this filtration is also presented in Figure S4E showing a clearer trend in activity time versus fluorescence and a flatter relationship between the mean *τ*_*OFF*_ and the mean fluorescence. We believe that these long periods are due to the *eve* regulation by multiple stripe-specific enhancers, in contrast to transgenic constructs, which typically correspond to the effects of a single enhancer. The complex interaction between these enhancers likely contributes to increased time variability while maintaining a similar spatial pattern of ⟨*τ*_*ON*_⟩ and ⟨*τ*_*OFF*_⟩ across stripes. In addition, it is known that *eve* stripe expression shifts anteriorly during NC14, such that some cells expressing *eve* at the posterior boundary of a stripe early in NC14 become transcriptionally inactive later [41]. For example, a cell located between stripe 1 and stripe 2 may initially express genes early in NC14 due to the activity of stripe 1 enhancers. Midway through NC14, it becomes transcriptionally inactive and later in NC14, as the expression domain shifts, it begins to express genes regulated by stripe 2-specific enhancers. This trajectory results in a long inactivity period between bursts, which contributes to higher variability. In fact, nuclei with inter-burst times of more than 12 min are mostly located at the edges of each stripe (Figure S4B).

In summary, our analysis suggests that the bursting properties of endogenous genes are comparable to those of developmental enhancers. These findings align with a previous study that characterized the bursting properties of *eve* using a reporter construct based on bacterial artificial chromosomes [48, 54]. Similarly to what has been observed in enhancers, the mean gene expression level for the *eve* gene is mainly aligned with changes in activity time despite showing a more complex spatial pattern (Figure 7A). We observed some cells with anomaly long interburst duration that biased the *τ*_*OFF*_ statistics. We hypothesized that these long periods are likely due to the coordinated action of multiple enhancers that collectively regulate *eve* expression.

## Discussion

In this article, we aim to explain how the spatial pattern of mean gene expression in the early embryonic development of *Drosophila* emerges from the temporal dynamics of transcriptional burst activity at the single cell level. We analyze the dynamics of the amount of nascent RNA molecules measured using MS2 tagging. To infer the transcriptional state (active or inactive), we propose an algorithm based on the signal slope (see Methods). The simplicity of our algorithm helps us analyze thousands of fluorescence trajectories from individual nuclei with high throughput, providing reproducible results on inferred transcriptional states. Our analysis reveals that fluorescence accumulation and decay follow linear functions of time (Figure S1). Although simple, this dynamics requires a careful explanation using a model that considers a detailed description of RNA polymerase recruitment and elongation on the gene. Although this kind of modeling is beyond the scope of this article, we show that the process can be simplified by a two-state model as described by equations (1) and (2) in Methods to capture key transcriptional burst properties.

On the basis of the statistics of transcriptional bursts, it is possible to infer properties of the mechanism controlling the burst timing. There are known mechanisms that drive and regulate transcriptional bursting, such as chromatin dynamics [55–59], 3D enhancer-promoter interactions [54, 60–63], transcription factor binding [64–68], mRNA polymerase pausing [69], and interactions between developmental promoter motifs and associated proteins [70]. Our observation of exponentially-distributed interburst timing (Figure 4B) suggests that burst initiation may be governed by a single rate-limiting step in these developmental enhancers. The observation of burst-duration timing to be hypoexponentially distributed as shown in Figure 4 B (that is, these times have statistical fluctuations less than an exponential distribution) implies a more complex transcriptional turn-OFF with multiple rate-limited steps. Findings of non-exponential promoter state durations have previously been made for other promoters in their OFF times [71, 72], and also for their ON times [73]. Further analysis of whether the durations of consecutive bursts are related (Figure S5) revealed weak correlations suggesting ON/OFF times being independent and identically distributed random variables as good approximations for the observed bursting behavior.

There is a rich body of literature connecting transcriptional bursting to fluctuations in levels of a given gene product [74–84]. Borrowing this recent mathematical modeling framework, we explore the impact of non-exponential promoter switching times on the statistical fluctuations in gene expression levels. We linked transcription states (ON/OFF) to the generation and degradation of a gene product, with the time spent in each state itself being an independent and identically distributed random variable following an arbitrary distribution (see Supplementary Information Section S5). Our analysis shows that the noise in mRNA levels increases with the noise in the time spent in either of the ON and OFF states (Figure S6). These results imply that tight control of transcriptional burst durations, as seen in this study, will function to buffer stochastic variations in expressed mRNA levels. An intriguing finding is the consistency in the statistics of promoter ON/OFF times across cells and different genes suggesting common mechanisms controlling bursting timing. It is worth mentioning that many studies have suggested that the kinetics of bursting can vary under different conditions [48, 85, 86], and that the time between successive bursts was observed to be controlled by the gene enhancer. This control includes, as shown here, decreasing the mean duration and regulating its random variability to hypoexponential levels. In future research, we plan to modify the distance and position of the enhancer relative to the promoter to better understand the mechanisms of bursting regulation.

We found that one of the main contributors to the formation of the gene expression pattern is activity time (Figure 5), and there is also a weak dependence of the mean ⟨*τ*_*OFF*_⟩ on transcription activity. This observation is similar to previous findings [85, 87]. The activity time and ⟨*τ*_*OFF*_⟩ are related variables and their regulation can share similar mechanisms of action. Specifically, we found cells with isolated periods of long inactivity (>12 min) in between bursts suggesting that these cells can have more than one activity time. In recent work [88], monitoring the transcriptional activity of a promoter under the control of Polycomb proteins has shown a switch between an OFF state and a transcriptionally *permissive* state, with multiple bursts occurring during the permissive period. In this context, the activity time can be understood as the permissive period that is developmentally regulated to drive spatial expression patterning.

The observed importance of activity time in controlling gene expression patterns raises interesting questions about its molecular regulation. Since activity time represents the temporal window during which a gene undergoes bursting cycles, it probably reflects the duration of allowed promoter states for transcriptional activation. We hypothesize that binding of transcription factors (TFs) to cis-regulatory sequences could control this window of activity. For example, Dorsal, a transcription factor that activates *sna* and *rho* studied here, shows a graded expression along the dorsoventral axis of the embryo, showing its highest expression at the most ventral point and a gradual decrease in expression level [89]. The activity time of the Dorsal target genes (*sna* and rho), correlates with Dorsal level. Specifically, Dorsal target genes show a longer activity time in the ventral region compared to the more dorsolateral domain. Moreover, cooperative binding of multiple TFs at enhancers may create a stable microenvironment that persists for the duration of the activity time. Our observation that different enhancers (such as *snaPE* versus *snaSE* ) exhibit distinct activity time distributions supports this model, as these enhancers contain different combinations of TF binding sites that could affect the stability and duration of active transcriptional states.

In summary, we present a simplified model to capture the stochastic dynamics of transcription in single cells and propose high-throughput techniques to infer promoter states. Our results reveal that these burst timing statistics exhibit uniform spatial patterns within the embryo, with ON times showing tighter statistical fluctuations compared to the OFF times. Moreover, a major determinant of spatial expression regulation for different developmental enhancers is the activity time that is related to the permissive period of transcription.

## Methods

### Experimental details

MS2/MCP system is used to visualize nascent transcripts (Figure 1C). To build the MS2/MCP mechanism, 24 copies of MS2 sequences are inserted into the 5’ UTR or 3’ UTR of the target gene of interest. Upon transcription, MS2 sequences form a stem-loop structure, which can bind to maternally loaded MCP proteins fused with GFP. As a result, clustered MCP-GFP molecules at the site of active transcription can be detected as fluorescent puncta in each nucleus (Figure 1B).

The data analyzed in this article, for *rho, Kr*, and *sna* genes, correspond to the experiments carried out by the reference [40], where the regulatory enhancer of each gene drives the expression of the *MS2-yellow* reporter gene (Figure 1C). For *eve*, we used data from [41], where endogenous *eve* is tagged with MS2 using CRISPR-mediated genome editing. For the nonendogenous data, it has been observed that the dynamics of the reporter gene recapitulate the expression of the endogenous gene. All images are taken in embryos 2-3 hours after fertilization, corresponding to the 14th nuclear cycle (NC14). From each live imaging movie, we track the fluorescence intensity of the MS2 puncta in each nucleus for the duration of NC14 and use it as a measure of instantaneous transcriptional activity (Figure 1D).

### Transcription state Inference

As shown in Figure 1D, a typical fluorescence trajectory reveals complex dynamics. We can segment the complete signal into periods of no activity when the fluorescence stays at a basal level (before the first burst and after the last one) and periods of activity when transcriptional bursting occurs. Our goal is to determine the precise moments at which these bursts occur which are related to the moment in which the fluorescence signal rises. To do so, we first explain the model on which the inference method is based and then explain the method itself.

#### Phenomenological modeling of the fluorescence signal dynamics

To simplify our analysis, we consider that the fluorescence signal dynamics is driven mainly by the transcriptional state. This state exists in two modes: ON and OFF. When the transcriptional state is ON, high transcriptional activity results in rising fluorescence levels (illustrated by dark green periods in Figure 1D). Conversely, during the OFF state, the absence of RNA polymerase recruitment causes the fluorescence signal to decrease or stabilize at its base level (light green periods in Figure 1D).

Given the possible transcriptional states (ON and OFF), we define the function *ϕ*(*t* ) as follows:

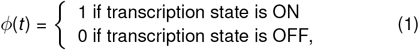

which quantifies the transcriptional state value. When the state is ON, we assume that the fluorescence levels increase at a constant rate. Similarly, when the gene is OFF, the fluorescence also decreases at a constant rate. This linear accumulation and decay approximation was concluded after a detailed analysis of fluorescence accumulation and decay trajectories in response to ON/OFF states, as presented in Figure S1. In addition to a good fit with linear functions of time, we observe that the slopes for decreasing/increasing dynamics have very similar mean absolute values (Figure S2).

In this context, we simplify the dynamics of the fluorescence signal assuming that the signal increases and decreases with the same absolute rate *λ* that could be different for each cell. Given the transcriptional state *ϕ*(*t* ), the fluorescence signal *y*^*m*^ is modeled as per the following ordinary differential equation:

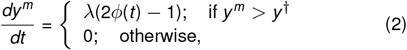

with the superscript *m* standing for the *model-predicted* signal. The burst threshold *y* ^†^ is defined such that below this threshold, we assume that no bursting occurs and the transcription state is OFF (Figure 1). To determine *y* ^†^, we selected the maximum fluorescence level in cells that did not exhibit noticeable bursts. This value was found to be *y* ^†^ = 0.1 (a.u.). Since basal fluorescence levels did not differ significantly between genes, we used the same threshold for all genes. Next, we will explain how, assuming (2), we infer the most probable *λ* and the corresponding transcriptional state for each cell.

#### Transcription state inference method

Since the data consists of sampling the fluorescence signal every Δ*t* = 20*s*, we consider a discrete-time analog of (2) that follows

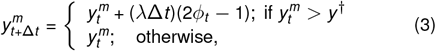

where *t* = {0, Δ*t* , 2Δ*t* , … }. Given experimental measurements *y*_*t*_ , our objective is to infer the transcriptional state *ϕ*_*t*_ that best explains the data. The technical details of the defined algorithm are presented in Supplementary Information Section S3, and are briefly summarized here.

The inference method starts with the experimental trajectory *y*_*t*_, which is first smoothed to obtain the signal 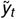 by convolving it with a triangular kernel function to not flatten the signal peaks too much (see Supplementary Information Section S3). Next, we propose a value for *λ* for all transcriptional bursts in a single nucleus. Given 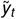 and *λ*, we first obtain the *transcription activity ρ*_*t*_ defined as

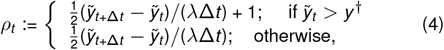

and is a quantity which estimates the rate of signal increase relative to *λ*. We perform a new round of smoothing of *ρ*_*t*_ with a square kernel function (see Supplementary Information section 3) obtaining the smoothed 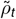. In this way, the inferred transcriptional state *ϕ*_*t*_ is calculated as follows:

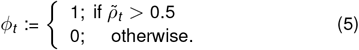

The resulting *ϕ*_*t*_ is replaced in (3) obtaining a trajectory 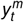 that depends parametrically on the particular *λ*. Finally, the optimal *λ*, denoted as *λ*^*^, is defined as the loading rate for that specific nuclei that minimizes the mean square distance between the data *y*_*t*_ and the resultant 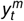 defined as:

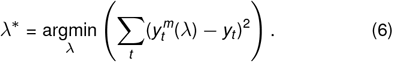

The optimal *λ*^*^ will predict the optimal *ϕ*_*t*_ by using (5) which is taken as the most accurate trajectory of the transcriptional state. Examples of best-fit trajectories *ϕ*_*t*_ and 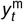 are shown in Figure S3D-E, respectively.

## Author Contributions

- **Conceptualization:** B.L. C.N., A.S.
- **Investigation:** Z.V., C.N.
- **Data Curation:** B.L., Z.V., C.N.
- **Formal Analysis:** Z.V., C.N.
- **Funding Acquisition:** A.S., B.L.
- **Investigation:** C.N., Z.V., B.L., A.S.
- **Methodology:** B.L., Z.V., C.N.
- **Resources:** A.S., B.L.
- **Software:** C.N., Z.V.
- **Supervision:** A.S., B.L.
- **Visualization:** C.N., Z.V.
- **Writing:** C.N., B.L., A.S.
- **Writing – Review Editing:** B.L., A.S.

## Competing interests

The authors have no conflict of interest to declare.

## Acknowledgments

B.L. is funded by NIH R35 GM133425. A.S. and C.N. acknowledge support from NIH-NIGMS through grant R35GM148351.

## Data accessibility statement

Data set and data processing scripts are publicly available at https://doi.org/10.5281/zenodo.13208281 [90].

## Notes

https://doi.org/10.5281/zenodo.13208281

### Competing Interest Statement

The authors have declared no competing interest.

## References

[1] Georg Halder, Patrick Callaerts, and Walter J Gehring. Induction of ectopic eyes by targeted expression of the eyeless gene in Drosophila. Science, 267(5205):1788–1792, 1995.

[2] Ingmar H Riedel-Kruse, Claudia Muller, and Andrew C Oates. Synchrony dynamics during initiation, failure, and rescue of the segmentation clock. Science, 317(5846):1911–1915, 2007.

[3] Julien O Dubuis, Reba Samanta, and Thomas Gregor. Accurate measurements of dynamics and reproducibility in small genetic networks. Molecular Systems Biology, 9(1):639, 2013.

[4] Noboru Jo Sakabe, Daniel Savic, and Marcelo A Nobrega. Transcriptional enhancers in development and disease. Genome Biology, 13(1):1–11, 2012.

[5] Annique Claringbould and Judith B Zaugg. Enhancers in disease: molecular basis and emerging treatment strategies. Trends in Molecular Medicine, 27(11):1060–1073, 2021.

[6] M. B. Elowitz, A. J. Levine, E. D. Siggia, and P. S. Swain. Stochastic gene expression in a single cell. Science, 297:1183–1186, 2002.

[7] Arjun Raj and Alexander Van Oudenaarden. Nature, nurture, or chance: stochastic gene expression and its consequences. Cell, 135(2):216–226, 2008.

[8] William R Holmes, Nabora Soledad Reyes de Mochel, Qixuan Wang, Huijing Du, Tao Peng, Michael Chiang, Olivier Cinquin, Ken Cho, and Qing Nie. Gene expression noise enhances robust organization of the early mammalian blastocyst. PLOS Computational Biology, 13(1):e1005320, 2017.

[9] Lukas Voortman and Robert J Johnston Jr. Transcriptional repression in stochastic gene expression, patterning, and cell fate specification. Developmental Biology, 481:129–138, 2022.

[10] Ilka Schultheiß. Araújo, Jessica Magdalena Pietsch, Emma Mathilde Keizer, Bettina Greese, Rachappa Balkunde, Christian Fleck, and Martin Hülskamp. Stochastic gene expression in arabidopsis thaliana. Nature Communications, 8(1):2132, 2017.

[11] Teresa Ferraro, Emilia Esposito, Laure Mancini, Sam Ng, Tanguy Lucas, Mathieu Coppey, Nathalie Dostatni, Aleksandra M Walczak, Michael Levine, and Mounia Lagha. Transcriptional memory in the Drosophila embryo. Current Biology, 26(2):212–218, 2016.

[12] Lei Zhang, Kelly Radtke, Likun Zheng, Anna Q Cai, Thomas F Schilling, and Qing Nie. Noise drives sharpening of gene expression boundaries in the zebrafish hindbrain. Molecular Systems Biology, 8(1):613, 2012.

[13] Ravi V Desai, Xinyue Chen, Benjamin Martin, Sonali Chaturvedi, Dong Woo Hwang, Weihan Li, Chen Yu, Sheng Ding, Matt Thomson, Robert H Singer, et al. A DNA repair pathway can regulate transcriptional noise to promote cell fate transitions. Science, 373(6557):eabc6506, 2021.

[14] Sevdenur Keskin, Gnanapackiam S Devakanmalai, Soo Bin Kwon, Ha T Vu, Qiyuan Hong, Yin Yeng Lee, Mohammad Soltani, Abhyudai Singh, Ahmet Ay, and Ertuğ rul M Özbudak. Noise in the vertebrate segmentation clock is boosted by time delays but tamed by notch signaling. Cell Reports, 23:2175–2185, 2018.

[15] Oriana QH Zinani, Kemal Keseroğ lu, Supravat Dey, Ahmet Ay, Abhyudai Singh, and Ertuğ rul M Özbudak. Gene copy number and negative feedback differentially regulate transcriptional variability of segmentation clock genes. iScience, 25(7), 2022.

[16] A. Raj, C.S. Peskin, D. Tranchina, D.Y. Vargas, and S. Tyagi. Stochastic mRNA synthesis in mammalian cells. PLOS Biology, 4:e309, 2006.

[17] Daniel Ramsköld, Gert-Jan Hendriks, Anton JM Larsson, Juliane V Mayr, Christoph Ziegenhain, Michael Hagemann-Jensen, Leonard Hartmanis, and Rickard Sandberg. Single-cell new RNA sequencing reveals principles of transcription at the resolution of individual bursts. Nature Cell Biology, 26(10):1725–1733, 2024.

[18] DM Jeziorska, EAJ Tunnacliffe, JM Brown, H Ayyub, J Sloane-Stanley, JA Sharpe, BC Lagerholm, C Babbs, AJH Smith, VJ Buckle, et al. On-microscope staging of live cells reveals changes in the dynamics of transcriptional bursting during differentiation. Nature Communications, 13(1):6641, 2022.

[19] Yihan Wan, Dimitrios G Anastasakis, Joseph Rodriguez, Murali Palangat, Prabhakar Gudla, George Zaki, Mayank Tandon, Gianluca Pegoraro, Carson C Chow, Markus Hafner, et al. Dynamic imaging of nascent RNA reveals general principles of transcription dynamics and stochastic splice site selection. Cell, 184(11):2878–2895, 2021.

[20] Emilia A Leyes Porello, Robert T Trudeau, and Bomyi Lim. Transcriptional bursting: stochasticity in deterministic development. Development, 150(12):dev201546, 2023.

[21] Takashi Fukaya. Enhancer dynamics: Unraveling the mechanism of transcriptional bursting. Science Advances, 9(31):eadj3366, 2023.

[22] Roy D. Dar, Sydney M. Shaffer, Abhyudai Singh, Brandon S. Razooky, Michael L. Simpson, Arjun Raj, and Leor S. Weinberger. Transcriptional bursting explains the noise–versus–mean relationship in mRNA and protein levels. PLOS ONE, 11:e0158298, 2016.

[23] Adam M. Corrigan and Jonathan R. Chubbemail. Regulation of transcriptional bursting by a naturally oscillating signal. Current Biology, 24:205–211, 2014.

[24] Hiroshi Ochiai, Tetsutaro Hayashi, Mana Umeda, Mika Yoshimura, Akihito Harada, Yukiko Shimizu, Kenta Nakano, Noriko Saitoh, Zhe Liu, Takashi Yamamoto, et al. Genome-wide kinetic properties of transcriptional bursting in mouse embryonic stem cells. Science Advances, 6:eaaz6699, 2020.

[25] Daniel R Larson, Christoph Fritzsch, Liang Sun, Xiuhau Meng, David S Lawrence, and Robert H Singer. Direct observation of frequency modulated transcription in single cells using light activation. eLife, 2:e00750, 2013.

[26] Massimo Cavallaro, Mark D Walsh, Matt Jones, James Teahan, Simone Tiberi, Bärbel Finkenstädt, and Daniel Heben-streit. 3’-5’ crosstalk contributes to transcriptional bursting. Genome Biology, 22:1–20, 2021.

[27] Sharon Yunger, Liat Rosenfeld, Yuval Garini, and Yaron Shav-Tal. Single-allele analysis of transcription kinetics in living mammalian cells. Nature Methods, 7(8):631–633, 2010.

[28] Daniel R Larson, Daniel Zenklusen, Bin Wu, Jeffrey A Chao, and Robert H Singer. Real-time observation of transcription initiation and elongation on an endogenous yeast gene. Science, 332(6028):475–478, 2011.

[29] Adam M Corrigan, Edward Tunnacliffe, Danielle Cannon, and Jonathan R Chubb. A continuum model of transcriptional bursting. eLife, 5:e13051, 2016.

[30] Jacques P Bothma, Hernan G Garcia, Emilia Esposito, Gavin Schlissel, Thomas Gregor, and Michael Levine. Dynamic regulation of eve stripe 2 expression reveals transcriptional bursts in living Drosophila embryos. Proceedings of the National Academy of Sciences, 111(29):10598–10603, 2014.

[31] Roy D Dar, Brandon S Razooky, Abhyudai Singh, Thomas V Trimeloni, James M McCollum, Chris D Cox, Michael L Simpson, and Leor S Weinberger. Transcriptional burst frequency and burst size are equally modulated across the human genome. Proceedings of the National Academy of Sciences, 109(43):17454–17459, 2012.

[32] Olivia Padovan-Merhar, Gautham P Nair, Andrew G Biaesch, Andreas Mayer, Steven Scarfone, Shawn W Foley, Angela R Wu, L Stirling Churchman, Abhyudai Singh, and Arjun Raj. Single mammalian cells compensate for differences in cellular volume and DNA copy number through independent global transcriptional mechanisms. Molecular Cell, 58(2):339–352, 2015.

[33] Shasha Chong, Chongyi Chen, Hao Ge, and X. Sunney Xie. Mechanism of transcriptional bursting in bacteria. Cell, 158:314–326, 2014.

[34] A. Singh, Brandon Razooky, Chris D. Cox, Michael L. Simpson, and Leor S. Weinberger. Transcriptional bursting from the HIV-1 promoter is a significant source of stochastic noise in HIV-1 gene expression. Biophysical Journal, 98:L32–L34, 2010.

[35] Lea Schuh, Michael Saint-Antoine, Eric M Sanford, Benjamin L Emert, Abhyudai Singh, Carsten Marr, Arjun Raj, and Yogesh Goyal. Gene networks with transcriptional bursting recapitulate rare transient coordinated high expression states in cancer. Cell Systems, 10:363–378, 2020.

[36] A. Singh, B. S. Razooky, R. D. Dar, and L. S. Weinberger. Dynamics of protein noise can distinguish between alternate sources of gene-expression variability. Molecular Systems Biology, 8:607, 2012.

[37] Jeff E Mold, Martin H Weissman, Michael Ratz, Michael Hagemann-Jensen, Joanna Hård, Carl-Johan Eriksson, Hosein Toosi, Joseph Berghenstråhle, Christoph Ziegenhain, Leonie von Berlin, et al. Clonally heritable gene expression imparts a layer of diversity within cell types. Cell Systems, 15(2):149–165, 2024.

[38] Jonathan R Chubb, Tatjana Trcek, Shailesh M Shenoy, and Robert H Singer. Transcriptional pulsing of a developmental gene. Current Biology, 16(10):1018–1025, 2006.

[39] Ido Golding, Johan Paulsson, Scott M Zawilski, and Edward C Cox. Real-time kinetics of gene activity in individual bacteria. Cell, 123(6):1025–1036, 2005.

[40] Takashi Fukaya, Bomyi Lim, and Michael Levine. Enhancer control of transcriptional bursting. Cell, 166(2):358–368, 2016.

[41] Bomyi Lim, Takashi Fukaya, Tyler Heist, and Michael Levine. Temporal dynamics of pair-rule stripes in living Drosophila embryos. Proceedings of the National Academy of Sciences, 115(33):8376–8381, 2018.

[42] Edouard Bertrand, Pascal Chartrand, Matthias Schaefer, Shailesh M Shenoy, Robert H Singer, and Roy M Long. Localization of ash1 mrna particles in living yeast. Molecular Cell, 2(4):437–445, 1998.

[43] Hernan G Garcia, Mikhail Tikhonov, Albert Lin, and Thomas Gregor. Quantitative imaging of transcription in living Drosophila embryos links polymerase activity to patterning. Current Biology, 23(21):2140–2145, 2013.

[44] Jächle Herbert, Michael Hoch, Michael J Ankratz, Nicole Gerwin, Frank Sauer, and Günter Brönner. Transcriptional control by Drosophila gap genes. Journal of Cell Science, 1992(Supplement_16):39–51, 1992.

[45] Y Tony Ip, Ronald E Park, David Kosman, Ethan Bier, and Michael Levine. The dorsal gradient morphogen regulates stripes of rhomboid expression in the presumptive neuroectoderm of the Drosophila embryo. Genes & Development, 6(9):1728–1739, 1992.

[46] Michael W Perry, Alistair N Boettiger, Jacques P Bothma, and Michael Levine. Shadow enhancers foster robustness of Drosophila gastrulation. Current Biology, 20(17):1562–1567, 2010.

[47] Tadaatsu Goto, Paul Macdonald, and Tom Maniatis. Early and late periodic patterns of even skipped expression are controlled by distinct regulatory elements that respond to different spatial cues. Cell, 57(3):413–422, 1989.

[48] Augusto Berrocal, Nicholas C Lammers, Hernan G Garcia, and Michael B Eisen. Kinetic sculpting of the seven stripes of the Drosophila even-skipped gene. eLife, 9:e61635, 2020.

[49] PW Ingham. The molecular genetics of embryonic pattern formation in drosophila. Nature, 335(6185):25–34, 1988.

[50] Michael Hoch, C Schröder, Eveline Seifert, and H Jäckle. cis-acting control elements for krüppel expression in the Drosophila embryo. The EMBO journal, 9(8):2587–2595, 1990.

[51] Dig B Mahat, Nathaniel D Tippens, Jorge D Martin-Rufino, Sean K Waterton, Jiayu Fu, Sarah E Blatt, and Phillip A Sharp. Single-cell nascent RNA sequencing unveils coordinated global transcription. Nature, pages 1–8, 2024.

[52] Jacques P Bothma, Hernan G Garcia, Samuel Ng, Michael W Perry, Thomas Gregor, and Michael Levine. Enhancer additivity and non-additivity are determined by enhancer strength in the Drosophila embryo. eLife, 4:e07956, 2015.

[53] Michael J Pankratz and Herbert Jäckle. Making stripes in the Drosophila embryo. Trends in Genetics, 6:287–292, 1990.

[54] Augusto Berrocal, Nicholas C Lammers, Hernan G Garcia, and Michael B Eisen. Unified bursting strategies in ectopic and endogenous even-skipped expression patterns. eLife, 12:RP88671, 2024.

[55] Michele Gabriele, Hugo B Brandão, Simon Grosse-Holz, Asmita Jha, Gina M Dailey, Claudia Cattoglio, Tsung-Han S Hsieh, Leonid Mirny, Christoph Zechner, and Anders S Hansen. Dynamics of ctcf-and cohesin-mediated chromatin looping revealed by live-cell imaging. Science, 376(6592):496–501, 2022.

[56] LaTasha CR Fraser, Ryan J Dikdan, Supravat Dey, Abhyudai Singh, and Sanjay Tyagi. Reduction in gene expression noise by targeted increase in accessibility at gene loci. Proceedings of the National Academy of Sciences, 118(42):e2018640118, 2021.

[57] Dennis May, Sangwon Yun, David G Gonzalez, Sangbum Park, Yanbo Chen, Elizabeth Lathrop, Biao Cai, Tianchi Xin, Hongyu Zhao, Siyuan Wang, et al. Live imaging reveals chromatin compaction transitions and dynamic transcriptional bursting during stem cell differentiation in vivo. eLife, 12:e83444, 2023.

[58] Charles A Kenworthy, Nayem Haque, Shu-Hao Liou, Panagiotis Chandris, Vincent Wong, Patrycja Dziuba, Luke D Lavis, Wei-Li Liu, Robert H Singer, and Robert A Coleman. Bromodomains regulate dynamic targeting of the pbaf chromatin-remodeling complex to chromatin hubs. Biophysical Journal, 121(9):1738–1752, 2022.

[59] Ineke Brouwer, Emma Kerklingh, Fred van Leeuwen, and Tineke L Lenstra. Dynamic epistasis analysis reveals how chromatin remodeling regulates transcriptional bursting. Nature structural & molecular biology, 30(5):692–702, 2023.

[60] Christopher H Bohrer and Daniel R Larson. Synthetic analysis of chromatin tracing and live-cell imaging indicates pervasive spatial coupling between genes. eLife, 12:e81861, 2023.

[61] Caroline R. Bartman, Sarah C. Hsu, Chris C.-S. Hsiung, Arjun Raj, and Gerd A. Blobel. Enhancer regulation of transcriptional bursting parameters revealed by forced chromatin looping. Molecular Cell, 62:237 –247, 2016.

[62] Irene Robles-Rebollo, Sergi Cuartero, Adria Canellas-Socias, Sarah Wells, Mohammad M Karimi, Elisabetta Mereu, Alexandra G Chivu, Holger Heyn, Chad Whilding, Dirk Dormann, et al. Cohesin couples transcriptional bursting probabilities of inducible enhancers and promoters. Nature Communications, 13(1):4342, 2022.

[63] Manyu Du, Simon Hendrik Stitzinger, Jan-Hendrik Spille, Won-Ki Cho, Choongman Lee, Mohammed Hijaz, Andrea Quintana, and Ibrahim I Cissé. Direct observation of a condensate effect on super-enhancer controlled gene bursting. Cell, 187(2):331–344, 2024.

[64] Samuel H Keller, Siddhartha G Jena, Yuji Yamazaki, and Bomyi Lim. Regulation of spatiotemporal limits of developmental gene expression via enhancer grammar. Proceedings of the National Academy of Sciences, 117(26):15096–15103, 2020.

[65] Sahla Syed, Yifei Duan, and Bomyi Lim. Modulation of protein-dna binding reveals mechanisms of spatiotemporal gene control in early Drosophila embryos. eLife, 12:e85997, 2023.

[66] Paula Dobrinić, Aleksander T Szczurek, and Robert J Klose. Prc1 drives polycomb-mediated gene repression by controlling transcription initiation and burst frequency. Nature Structural & Molecular Biology, 28(10):811–824, 2021.

[67] Liang Ma, Zeyue Gao, Jiegen Wu, Bijunyao Zhong, Yuchen Xie, Wen Huang, and Yihan Lin. Co-condensation between transcription factor and coactivator p300 modulates transcriptional bursting kinetics. Molecular Cell, 81(8):1682–1697, 2021.

[68] Charis Fountas and Tineke L Lenstra. Better together: how cooperativity influences transcriptional bursting. Current Opinion in Genetics & Development, 89:102274, 2024.

[69] Katjana Tantale, Encar Garcia-Oliver, Marie-Cécile Robert, Adèle LÕhostis, Yueyuxiao Yang, Nikolay Tsanov, Rachel Topno, Thierry Gostan, Alja Kozulic-Pirher, Meenakshi Basu-Shrivastava, et al. Stochastic pausing at latent hiv-1 promoters generates transcriptional bursting. Nature Communications, 12(1):4503, 2021.

[70] Virginia L Pimmett, Matthieu Dejean, Carola Fernandez, Antonio Trullo, Edouard Bertrand, Ovidiu Radulescu, and Mounia Lagha. Quantitative imaging of transcription in living Drosophila embryos reveals the impact of core promoter motifs on promoter state dynamics. Nature Communications, 12(1):4504, 2021.

[71] David M Suter, Nacho Molina, David Gatfield, Kim Schneider, Ueli Schibler, and Felix Naef. Mammalian genes are transcribed with widely different bursting kinetics. Science, 332(6028):472–474, 2011.

[72] Sandeep Choubey, Jane Kondev, and Alvaro Sanchez. Deciphering transcriptional dynamics in vivo by counting nascent RNA molecules. PLOS Computational Biology, 11(11):e1004345, 2015.

[73] Katjana Tantale, Florian Mueller, Alja Kozulic-Pirher, Annick Lesne, Jean-Marc Victor, Marie-Cécile Robert, Serena Capozi, Racha Chouaib, Volker Bäcker, Julio Mateos-Langerak, et al. A single-molecule view of transcription reveals convoys of RNA polymerases and multi-scale bursting. Nature Communications, 7(1):12248, 2016.

[74] Zikai Xu, Mohammad Soltani, and Abhyudai Singh. Exact statistical moments of multi-mode stochastic hybrid systems with renewal transitions. In 2018 IEEE Conference on Decision and Control (CDC), pages 3510–3515. IEEE, 2018.

[75] Tao Jia and Rahul V. Kulkarni. Intrinsic noise in stochastic models of gene expression with molecular memory and bursting. Journal of Mathematical Biology, 106:058102, 2011.

[76] Renjie Wu, Bangyan Zhou, Wei Wang, and Feng Liu. Regulatory mechanisms for transcriptional bursting revealed by an event-based model. Research, 6:0253, 2023.

[77] Changhong Shi, Xiyan Yang, Jiajun Zhang, and Tianshou Zhou. Stochastic modeling of the mrna life process: A generalized master equation. Biophysical Journal, 122(20):4023–4041, 2023.

[78] Changhong Shi, Xiyan Yang, Tianshou Zhou, and Jiajun Zhang. Nascent RNA kinetics with complex promoter architecture: Analytic results and parameter inference. Physical Review E, 110(3):034413, 2024.

[79] Meiling Chen, Songhao Luo, Mengfang Cao, Chengjun Guo, Tianshou Zhou, and Jiajun Zhang. Exact distributions for stochastic gene expression models with arbitrary promoter architecture and translational bursting. Physical Review E, 105(1):014405, 2022.

[80] Adam R Stinchcombe, Charles S Peskin, and Daniel Tranchina. Population density approach for discrete mrna distributions in generalized switching models for stochastic gene expression. Physical Review E—Statistical, Nonlinear, and Soft Matter Physics, 85(6):061919, 2012.

[81] Niraj Kumar, Abhyudai Singh, and Rahul V. Kulkarni. Transcriptional bursting in gene expression: analytical results for general stochastic models. PLOS Computational Biology, 11:e1004292, 2015.

[82] Zahra Vahdat and Abhyudai Singh. Quantifying statistics of gene product copy-number fluctuations: A stochastic hybrid systems approach. In 2024 IEEE 63rd Conference on Decision and Control (CDC), pages 7792–7797, 2024.

[83] Zhixing Cao, Tatiana Filatova, Diego A Oyarzún, and Ramon Grima. A stochastic model of gene expression with polymerase recruitment and pause release. Biophysical Journal, 119(5):1002–1014, 2020.

[84] A Singh. Transient changes in intercellular protein variability identify sources of noise in gene expression. Biophysical Journal, 107:2214–2220, 2014.

[85] Benjamin Zoller, Shawn C Little, and Thomas Gregor. Diverse spatial expression patterns emerge from unified kinetics of transcriptional bursting. Cell, 175(3):835–847, 2018.

[86] Lauren Forbes Beadle, Hongpeng Zhou, Magnus Rattray, and Hilary L Ashe. Modulation of transcription burst amplitude underpins dosage compensation in the Drosophila embryo. Cell Reports, 42(4), 2023.

[87] Nacho Molina, David M Suter, Rosamaria Cannavo, Benjamin Zoller, Ivana Gotic, and Félix Naef. Stimulus-induced modulation of transcriptional bursting in a single mammalian gene. Proceedings of the National Academy of Sciences, 110(51):20563–20568, 2013.

[88] Aleksander T Szczurek, Emilia Dimitrova, Jessica R Kelley, Neil P Blackledge, and Robert J Klose. The polycomb system sustains promoters in a deep off state by limiting pre-initiation complex formation to counteract transcription. Nature Cell Biology, 26(10):1700–1711, 2024.

[89] Joung-Woo Hong, David A Hendrix, Dmitri Papatsenko, and Michael S Levine. How the dorsal gradient works: insights from postgenome technologies. Proceedings of the National Academy of Sciences, 105(51):20072–20076, 2008.

[90] Cesar Nieto and Zahra Vahdat. Data processing pipeline for the article “Analysis of Transcriptional Bursting Dynamics and its Role on Spatial Patterning of D. melanogaster Embryo development” doi:10.5281/zenodo.13324127, August 2024.

