## Supplementary Information for "Regulation of Transcriptional Bursting and Spatial Patterning in Early *Drosophila* Embryo Development"

### S1 Analysis of single trajectories of fluorescence accumulation and decay

We first characterize the dynamics of fluorescence accumulation and decay as the promoter transitions between transcriptionally active and inactive states, respectively. Depending on how far these processes occur from saturation, one could expect these trajectories to be either exponential functions of time or well approximated by linear functions of time. To discriminate between these hypotheses, we plot the dynamics of fluorescence accumulation and decay for some long-duration bursts from one replicate of *Kr* gene in Figure S1. We also compare the data and the fit to linear functions of time (solid lines). Given the relatively good alignment of the data with linear functions of time, we propose a simplified model for the fluorescence intensity dynamics in response to promoter-state fluctuations as given by (3).

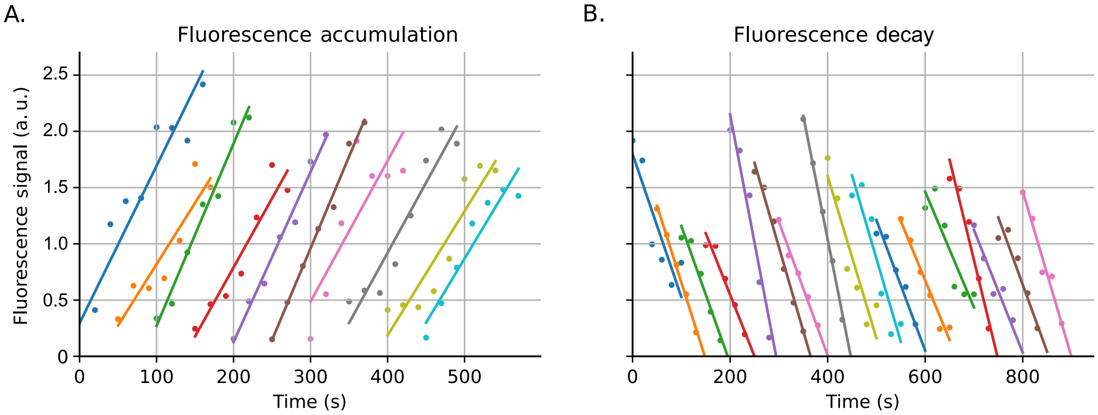


Figure S1: **Linear fit of fluorescence accumulation and decay trajectories over time.** Different colors represent fluorescence trajectories over time with their respective linear fit. (**A**) Fluorescence accumulation. (**B**) Fluorescence decay. For this plot, we selected random fluorescence trajectories with at least seven data points for accumulation and five data points for decrease were analyzed. These trajectories are selected from Replicate 1 the *Kr* gene.

### S2 Statistics of fluorescence accumulation and decay rates across genetic constructs

We aim to understand how the rate of fluorescence accumulation (related to RNA polymerase loading rate) varies for individual burst events in the same cell, across cells, and across genes. Moreover, how different are the accumulation rates from the decay rates? Thus, we combine the slopes of the linear fitted functions (Figure S1) for all the trajectories of fluorescence accumulation and decay, as shown in Figure S1. The resulting histograms for the absolute values of these slopes are presented in Figure S2. We observe that the mean rates have relatively low variability across different strains. In addition, the values of the decay rates are statistically similar to the accumulation rates. These findings motivate us to approximate the accumulation rate with the decay rate in our transcriptional dynamics model (3), significantly simplifying the inference algorithm.

### S3 Details of the applied inference algorithm

The previous analyses show the validity of the strongest simplifications made in our model. Fluorescence accumulation and decay were taken as linear functions of time and assumed to be approximately the same value that can vary from cell to cell. Here, we show three examples of fitting the experimental data and the inferred promoter states by applying our inference algorithm (Figure S3). Specifically, our algorithm consists of the following steps:

- **Defining a burst threshold**: When the fluorescence signal is above the burst threshold level $y^{\dagger}$ can be considered a burst

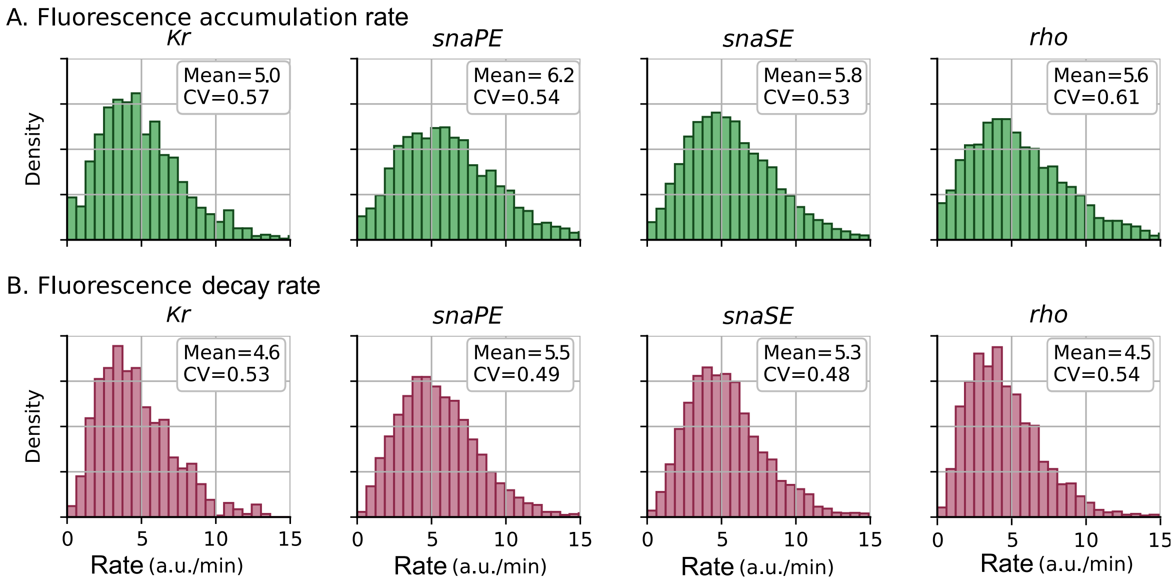


Figure S2: **Statistics of fluorescence accumulation and decay rates.** Distribution of the best slope of different trajectories of accumulation (**A**) and Fluorescence decay (**B**). To obtain the histograms, we fit trajectories of accumulation and decay of burst with durations longer than 1 minute. To obtain each histogram, data from the three studied replicas for each gene were pooled. The average mean slope and its coefficient of variation are shown in the inset.

(Figure S3A (dashed line)). To define the value of $y^{\dagger}$, we select cells without visible peaks (not shown in the plot). The threshold is defined as the average of the maximum points of these fluorescence trajectories. Given that this threshold was approximately consistent between strains, we set a fixed value which, in the units presented in the article, is $y^{\dagger} = 0.1$ a.u.

- **Signal smoothing**: Smoothing is performed by convolving the data with a 7-point triangular kernel. This smoothing follows the formula:

$$\tilde{y}_t = \sum_{t'=-d}^{d} K_{t'}^{d} y_{t-t'} \tag{S1}$$

in which $K_{t'}^{d}$ is the kernel. For the triangular kernel:

$$K_{t'}^{d} = \left(1 - \left|\frac{t'}{d}\right|\right); \qquad -d < t' < d \tag{S2}$$

the $d = 7$ is the length of the kernel. The idea of using this kernel is to not truncate the peaks of the signal (Figure S3B).

- **Inferring the transcriptional activity** $\rho_t$: We apply (4) to the smoothed signal in the previous step (Figure S3C).

- **Smoothing** $\rho_t$: We perform a convolution of the resultant $\rho_t$ with a square kernel with $d = 2$ points. This convolution was performed using the square kernel with the formula

$$\tilde{\rho}_t = \sum_{t'=-d}^{d} K_{t'}^{d} \rho_{t-t'} : \text{with } K_{t'}^{d} = \frac{1}{2d}; \qquad -d < t' < d \tag{S3}$$

The application of this function is to smooth the peaks in $\rho_t$ and to buffer the random fluctuations of the signal (Figure S3C).

- **Transcriptional state inference**: After obtaining $\tilde{\rho}_t$, we apply (5) to obtain the transcriptional state $\phi_t$. Examples of the results obtained after this step are shown in Figure S3D.

- **Comparison of the inferred** $y_t^m$ **with the original signal** $y_t$: Inferred $\phi_t$ is used in formula (3) to obtain a predicted signal $y_t^m$. The most probable inferred gene state is the one that minimizes the mean squared distance between $y_t^m$ and $y_t$. Examples of the most probable $y_t^m$ are shown in Figure S3E.

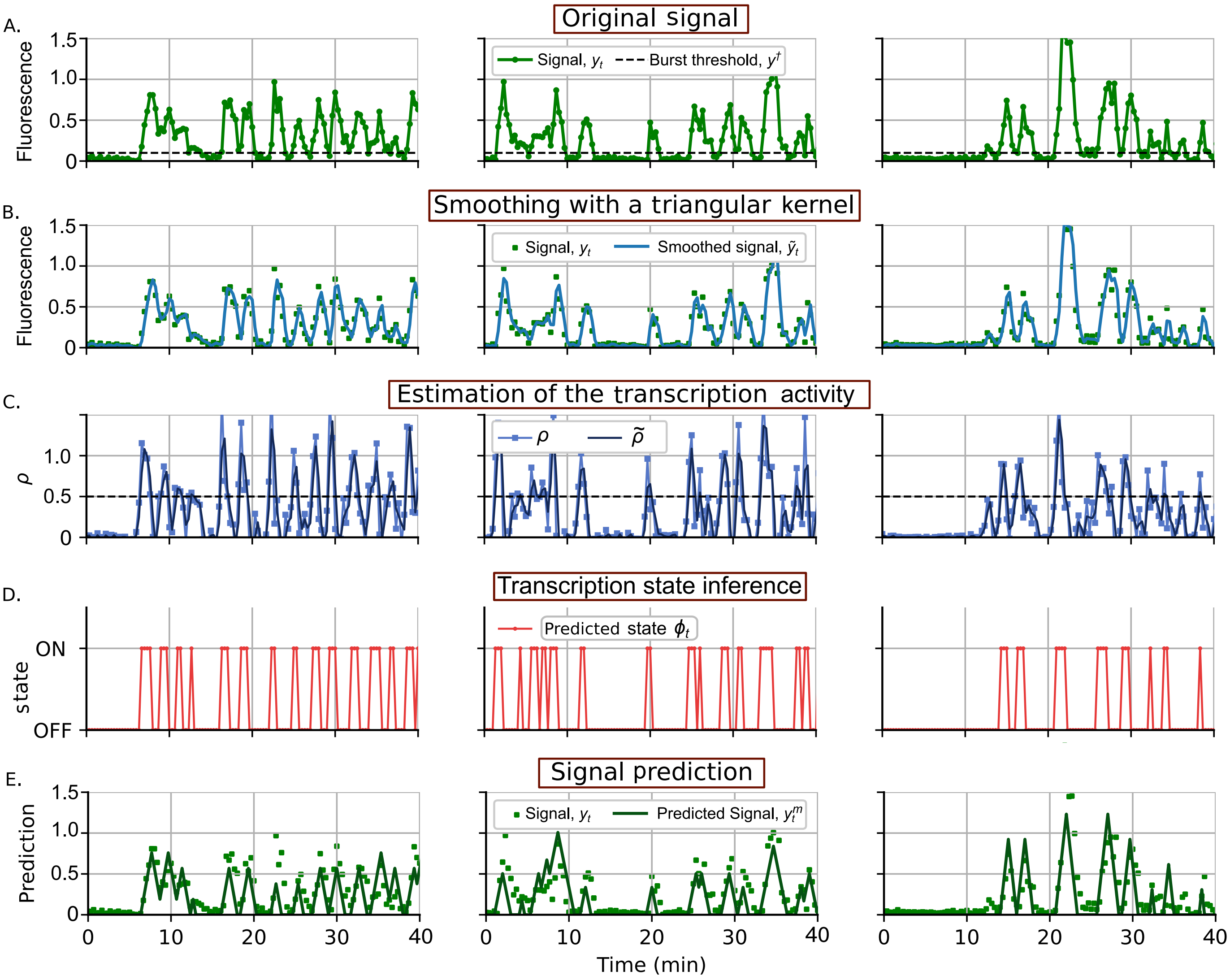


Figure S3: **Inference process applied to three different trajectories of single-cell fluorescence intensity.** (**A**) Original fluorescence over time $y_t$ in a single cell (see experimental method in Figure 1). Black dashed line represents the burst threshold. (**B**) Signal $\tilde{y}_t$ obtained after smoothing through convolution with a triangular kernel (S2). (**C**) Obtained transcription activity $\rho$ using (4) compared with the smoothed $\tilde{\rho}_t$ after convolution with a square kernel (S3). The dashed line shows that when $\tilde{\rho}_t > 0.5$, the gene state is considered as ON. (**D**) Inferred gene state using (5). (**E**) (solid line) Predicted fluorescence $y_t^m$ trajectory using the inferred transcriptional state $\phi_t$ and the optimal loading rate $\lambda^*$. This inferred signal is compared with the original signal $y_t$ (dots).

### S4 Correlation of timing properties for consecutive bursts events

In Figure S5, we present the results of the correlation of timing properties for consecutive bursts events for three replicas of each of the studied enhancers (*Kr*, *snaPE*, *snaSE*, *rho* and *eve*). We derive the ON and OFF intervals from the outcomes of the inference algorithm applied to the transcriptional trajectories of the five transgenic constructs. These intervals represent the periods when transcription is active (ON) and inactive (OFF), respectively. Figure S5A, represents the correlation between two consecutive ON intervals for the same transcription trajectory. Figure S5B illustrates the correlation between two consecutive "OFF" intervals. Figure S5C displays the correlation between the "ON" intervals and their consecutive "OFF" intervals. Lastly, Figure S5D shows the correlation between the "OFF" intervals and their consecutive "ON" intervals.

### S5 Gene expression noise with non-exponential switching between promoter states

In this section we study how stochastic switching between promoter states propagates to drive fluctuations in the mRNA levels. To focus primarily on the promoter fluctuations we consider deterministic time evolution of mRNA levels $x(t)$

$$\frac{dx}{dt} = \begin{cases} k - \gamma x & ;\text{ if gene is ON} \\ -\gamma x & ;\text{if gene is OFF,} \end{cases} \tag{S4}$$

which means that when the gene is ON, mRNAs are produced at a constant rate $k$. When the gene is OFF, each mRNA degrades with rate $\gamma$. We solve the moments in steady-state $\langle x \rangle$ and $\langle x^2 \rangle$ where $\langle\ \rangle$ represents the steady-state expected value. We refer the

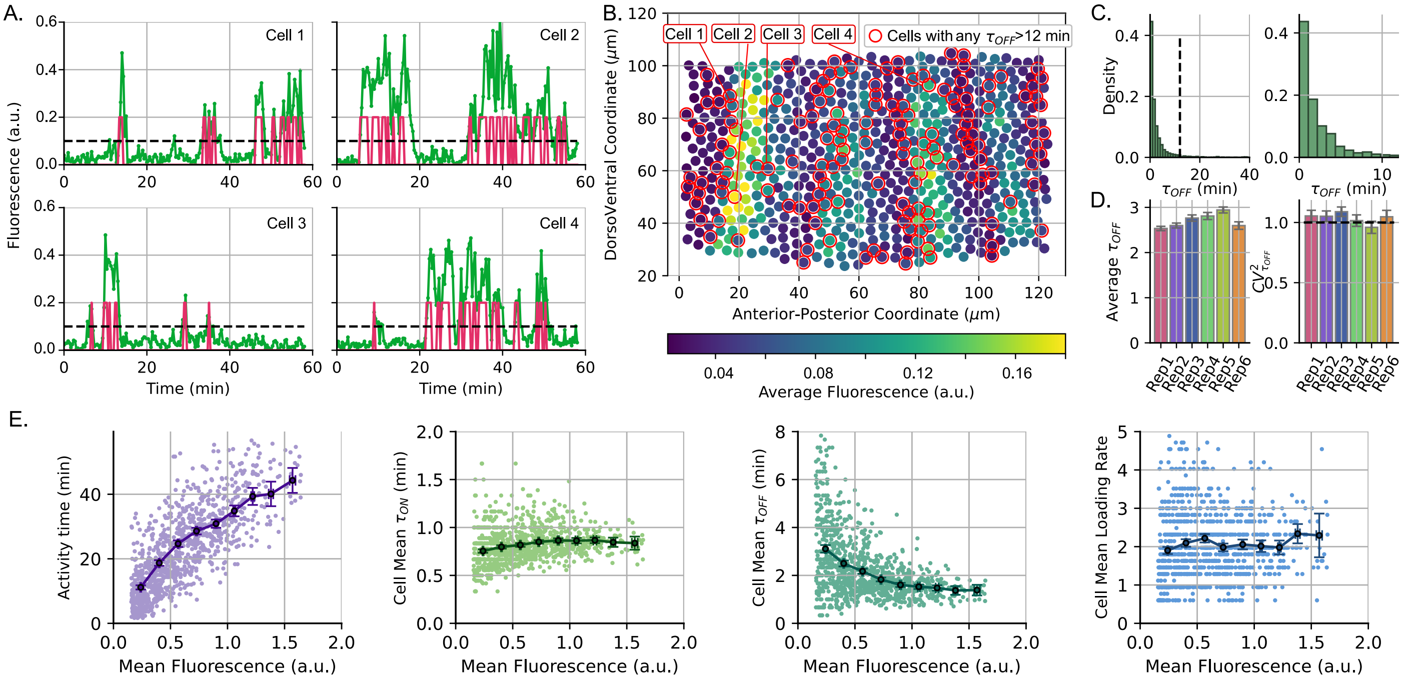


Figure S4: **Low transcription activity is related to the occurrence of unusually long periods of burst inactivity $\tau_{OFF}$ in the endogenous *eve* gene: (A)** Example of fluorescence trajectories with at least one $\tau_{OFF}$ exceeding 12 minutes. The green line represents the observed data, the red line indicates the inferred gene state, and the dashed black line marks the burst threshold. **(B)** Average fluorescence levels in the *eve* gene in replica 3. Each dot represents a single cell with coordinates estimated in the beginning of NC14. The dots are colored according to the cell average fluorescence level. Cells with any $\tau_{OFF}$ exceeding 12 minutes are highlighted. The plot suggests that these cells tend to be located in the edges of the *eve* stripes. **(C)** Distribution of all $\tau_{OFF}$ durations: *Left:* Distribution for all cells. *Right:* Distribution after excluding cells with at least one $\tau_{OFF}$ exceeding 12 minutes. **(D)** *Left:* Average $\tau_{OFF}$ after filtering out cells with any $\tau_{OFF}$ exceeding 12 minutes. *Right:* Variability in the time between bursts ($CV^2_{\tau_{OFF}}$) excluding inter-burst intervals with $\tau_{OFF}$ exceeding 12 minutes. Horizontal dashed line indicates the variability of the exponential distribution. Error bars show the 95% confidence interval of the estimated statistics using bootstrapping methods. **(E)** Trends of activity time, cell mean $\tau_{ON}$, cell mean $\tau_{OFF}$ and loading rate versus cell mean fluorescence after removing cells with any $\tau_{OFF} > 12$ min.

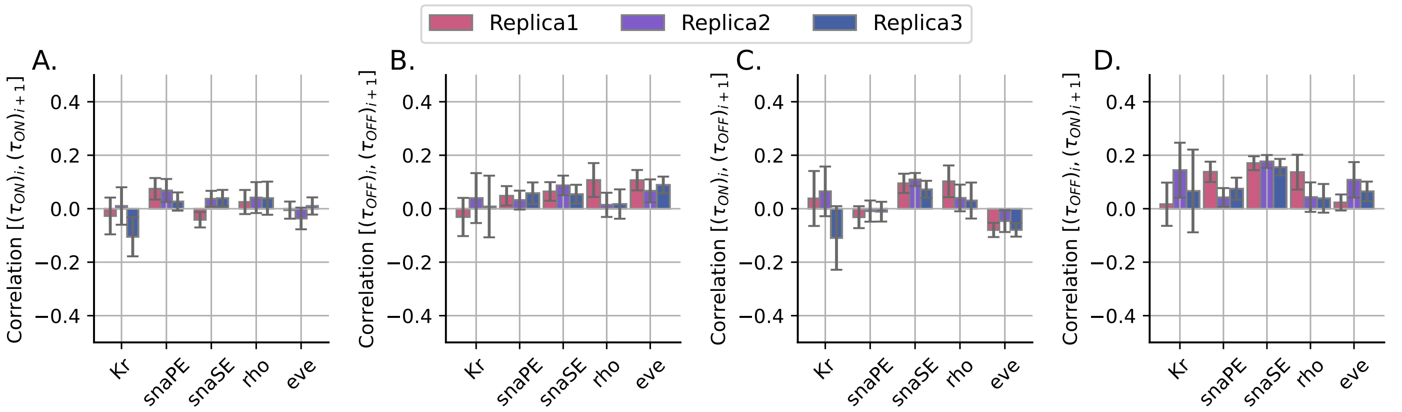


Figure S5: **Correlation of timing properties for consecutive bursts events** (**A**) Correlation between the duration of two consecutive bursts (**B**) Correlation between two consecutive "OFF" intervals (**C**) Correlation between the "ON" intervals and their consecutive "OFF" intervals. (**D**) Correlation between the "OFF" intervals and their consecutive "ON" intervals. Each color bar represents one replica of the studied genes (*Kr*, *snaPE*, *snaSE*, *rho* and *eve*). Three replicas in total for each gene are presented. Error-bars represent the 95% confidence interval using bootstrapping methods.

reader to [1, 2] for detailed derivations and similar approaches have been taken in other works [3–7]. The steady-state mean and the

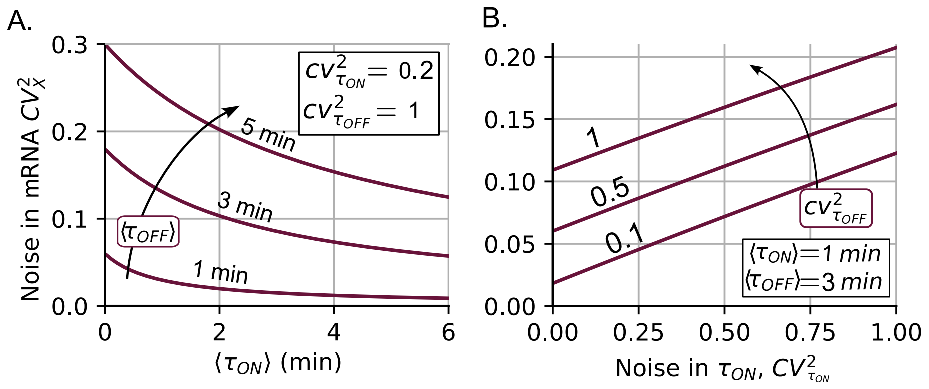


Figure S6: **Noise in mRNA levels considering non-exponentially distributed promoter switching times.** (**A**) Assuming gamma-distributed promoter switching times, $CV_x^2$ from (S7) is plotted as a function of $\langle\tau_{ON}\rangle$ for different values of $\langle\tau_{OFF}\rangle$ for $CV^2_{\tau_{ON}}$ = 0.2 and $CV^2_{\tau_{ON}}$ = 1. (**B**) $CV_x^2$ as a function of $CV^2_{\tau_{ON}}$ for different values of $CV^2_{\tau_{OFF}}$ using as parameters $\langle\tau_{ON}\rangle$ = 1 min, $\langle\tau_{OFF}\rangle$ = 3 min, and mRNA degradation rate is $\gamma = \ln(2)/7\,\text{min}^{-1}$ and the mRNA mean level is fixed at 20 as per (S5) implying $k = 7.92\,\text{min}^{-1}$.

noise (the square of the coefficient of variation) in mRNA levels are derived as:

$$\langle x\rangle = \frac{k}{\gamma}\frac{1}{1+\frac{\langle\tau_{OFF}\rangle}{\langle\tau_{ON}\rangle}} \tag{S5}$$

$$CV_x^2 = \frac{\left(1-\langle e^{-\gamma\tau_{OFF}}\rangle\right)\left(1-\langle e^{-\gamma\tau_{ON}}\rangle\right)}{\left(\langle e^{-\gamma\tau_{OFF}}\rangle\langle e^{-\gamma\tau_{ON}}\rangle-1\right)}\frac{1}{\gamma\langle\tau_{ON}\rangle}\left(\frac{\langle\tau_{OFF}\rangle}{\langle\tau_{ON}\rangle}+1\right)+\frac{\langle\tau_{OFF}\rangle}{\langle\tau_{ON}\rangle}, \tag{S6}$$

respectively. Note that the model assumes the $\tau_{ON}$ and $\tau_{OFF}$ are independent and identically distributed variables following any arbitrary positive-valued distribution. In the specific case where they follow the gamma distribution with means ($\langle\tau_{ON}\rangle$, $\langle\tau_{OFF}\rangle$) and noises ($CV^2_{\tau_{ON}}$, $CV^2_{\tau_{OFF}}$), respectively, the noise of the gene product can be expressed as:

$$CV_x^2 = \left(\frac{1}{1-\left(CV^2_{\tau_{OFF}}\gamma\langle\tau_{OFF}\rangle+1\right)^{1/CV^2_{\tau_{OFF}}}}+\frac{1}{1-\left(CV^2_{\tau_{ON}}\gamma\langle\tau_{ON}\rangle+1\right)^{1/CV^2_{\tau_{ON}}}}-1\right)^{-1}\frac{1}{\gamma\langle\tau_{ON}\rangle}\left(\frac{\langle\tau_{OFF}\rangle}{\langle\tau_{ON}\rangle}+1\right)+\frac{\langle\tau_{OFF}\rangle}{\langle\tau_{ON}\rangle}. \tag{S7}$$

Figure S6A illustrates that the noise decreases with $\langle\tau_{ON}\rangle$ and increases with $\langle\tau_{OFF}\rangle$. In Figure S6B, the noise term $CV_x^2$ is an increasing function of both $CV^2_{\tau_{ON}}$ and $CV^2_{\tau_{OFF}}$, given the observed values of $\langle\tau_{ON}\rangle$ and $\langle\tau_{OFF}\rangle$.

### References


[1] Zahra Vahdat and Abhyudai Singh. Quantifying statistics of gene product copy-number fluctuations: A stochastic hybrid systems approach. In *2024 IEEE 63rd Conference on Decision and Control (CDC)*, pages 7792–7797, 2024.

[2] Zikai Xu, Mohammad Soltani, and Abhyudai Singh. Exact statistical moments of multi-mode stochastic hybrid systems with renewal transitions. In *2018 IEEE Conference on Decision and Control (CDC)*, pages 3510–3515. IEEE, 2018.

[3] Changhong Shi, Xiyan Yang, Jiajun Zhang, and Tianshou Zhou. Stochastic modeling of the mrna life process: A generalized master equation. *Biophysical Journal*, 122(20):4023–4041, 2023.

[4] Changhong Shi, Xiyan Yang, Tianshou Zhou, and Jiajun Zhang. Nascent RNA kinetics with complex promoter architecture: Analytic results and parameter inference. *Physical Review E*, 110(3):034413, 2024.

[5] Meiling Chen, Songhao Luo, Mengfang Cao, Chengjun Guo, Tianshou Zhou, and Jiajun Zhang. Exact distributions for stochastic gene expression models with arbitrary promoter architecture and translational bursting. *Physical Review E*, 105(1):014405, 2022.

[6] Adam R Stinchcombe, Charles S Peskin, and Daniel Tranchina. Population density approach for discrete mrna distributions in generalized switching models for stochastic gene expression. *Physical Review E—Statistical, Nonlinear, and Soft Matter Physics*, 85(6):061919, 2012.

[7] Niraj Kumar, Abhyudai Singh, and Rahul V. Kulkarni. Transcriptional bursting in gene expression: analytical results for general stochastic models. *PLOS Computational Biology*, 11:e1004292, 2015.
